# Stretching the Limits: From Planar-Biaxial Stress–Stretch to Arterial Pressure–Diameter

**DOI:** 10.1101/2025.07.17.665394

**Authors:** Thibault Vervenne, Nic Vermeeren, Nele Demeersseman, Heleen Fehervary, Mathias Peirlinck, Ellen Kuhl, Nele Famaey

## Abstract

Understanding the physiological condition of the vascular system is critical to explain, treat, and manage vascular disease. Numerous experimental and computational studies characterize the mechanical behavior of arterial tissue under controlled laboratory conditions. However, translating this knowledge into physiologically realistic conditions remains challenging. Key difficulties include selecting suitable and relevant test methods, minimizing uncertainty, and ensuring robust model validation. We present a novel integrative approach to translate laboratory experiments on arterial samples into clinically relevant pressure–diameter behavior. We perform controlled planar-biaxial tests on carotid arteries under three stretch ratios and generate axial and circumferential stress–stretch data to calibrate a fiber-reinforced soft tissue model. Using an analytical thick-walled cylindrical model, we predict subject-specific pressure–diameter behavior, informed by arterial prestretches from ring opening experiments. We systematically compare predictions against extension-inflation experiments on tubes from the same artery by applying controlled pairs of axial stretch and inner pressure, while recording outer diameter. We quantify prediction error in absolute and relative stretch regimes and evaluate the importance of the load-free reference dimensions. Results show how planar-biaxial tests probe different stretch regimes compared to extension-inflation deformations, leading to extrapolation of model predictions. We demonstrate how the constitutive material parameters can be fitted to different biomechanical loading conditions and assess the sensitivity of the simulations to axial stretch and circumferential prestretch. Only when key model parameters are accurately captured and their uncertainty propagated, planar-biaxial stress–stretch data can reliably predict arterial pressure–diameter behavior.

## 1. Introduction

Biological tissues exhibit complex structural, compositional and mechanical behavior, making it challenging to accurately quantify their biomechanical properties. Inverse modeling informed by experimental data offers a valuable approach for estimating these properties, but its reliability depends on careful selection of the appropriate test method, design of a physiologically relevant test protocol, making appropriate model assumptions, and performing robust validation. Factors such as biological variability, nonlinearity, and anisotropy further challenge reliable evaluation. This work focuses on key challenges in mechanical testing and subsequent parameter estimation for the biomechanics of arteries, aiming to contribute to more robust and interpretable material descriptions.

Planar-biaxial testing provides a unique and physiologically relevant method for characterizing the mechanical response of arterial tissue, under controlled multi-axial loading [1]. Unlike uniaxial tests, biaxial loading replicates the simultaneous circumferential and axial stresses experienced by arteries *in vivo*, offering a more complete picture of wall mechanics. A key advantage of this approach is also its tissue-sparing nature, with only small, flat specimens required [2].

While planar-biaxial testing provides detailed insight into the material behavior, it does not directly result in a pressure– diameter (*P*–*d*) relationship, which remains the clinically most relevant descriptor of arterial compliance and function. Pressure–diameter is especially important for cardiovascular tissue under dynamic loading conditions, such as those observed between systolic and diastolic blood pressures. *In vivo*, this pulsatility can be observed through ultrasound, MRI, or 4D-CT imaging [3, 4]. *Ex vivo* pressure–diameter can be captured through extension-inflation testing. Especially for small vessels, planar-biaxial tests are more challenging, and pressure–diameter tests are more intuitive from a mechano-clinical point of view [5]. Nevertheless, extension-inflations also suffer from the need of larger tissue sample dimensions and specific testing device requirements.

Stress–stretch data in the axial and circumferential direction is often used to identify suitable constitutive models describing the mechanical behavior of the relevant cardiovascular tissue. Depending on the symmetry and directional dependence observed in the material response, these models may assume isotropic or anisotropic characteristics. From the experimental stress–stretch data, specific material parameters can be estimated through model fitting, allowing the strain energy function to capture the nonlinear mechanical behavior of the tissue. Once calibrated, this constitutive model can be implemented in numerical simulations to predict tissue deformation under physiological conditions. In particular, it enables the computation of pressure–diameter (*P*–*d*) responses under simulated extension-inflation scenarios, effectively linking experimental mechanics with computer models [6, 7].

Despite the potential to derive clinically relevant insights from mechanical testing and computational modeling, researchers currently face several important challenges throughout this process. First, selecting the most appropriate mechanical testing method, whether planar-biaxial or extension-inflation, is not straightforward. The choice of the experiment depends on the specific output goals, tissue sample, and infrastructure availability. Second, designing a relevant experimental protocol is critical: planar-biaxial testing requires careful control of the axial and circumferential loading paths, while extension-inflation testing invokes simultaneous axial loading and pressurization, a closer approximation of *in vivo* conditions. Third, minimizing measurement uncertainty is essential, as errors in estimating stress (often due to geometric approximations such as wall thickness and cross-sectional area) and stretch (typically captured via digital image correlation or marker tracking) can significantly affect model calibration. Fourth, selecting appropriate modeling assumptions, including the choice of constitutive model and the boundary conditions applied in simulations, requires a balance between physiological realism and computational tractability. Specifically, axial and circumferential prestretch can be highly sensitive to modeled pressure–diameter outputs. Finally, robust model validation against independent experimental data is necessary to ensure predictive reliability, while remaining a non-trivial step due to variability in biological samples and limited available data. Each of these aspects demands careful consideration to ensure that experimental and computational findings meaningfully reflect true *in vivo* tissue behavior. In this study, we critically evaluate the results of planar-biaxial tensile testing and compare them against the results of extension-inflation tests, aiming to quantify the effects of measurement uncertainty and experimental and modeling design choices.

## 2. Materials and Methods

Fig. 1 schematically illustrates the integrated experimental and computational framework. All mechanical tests and measurements were carried out at FIBEr, the KU Leuven Core Facility for Biomechanical Experimentation. The resulting datasets span multiple mechanical tests, processing workflows, and modeling approaches. To promote transparency, reproducibility, and further research, all protocols, raw data, and post-processing code have been made publicly available.

**Figure 1:**
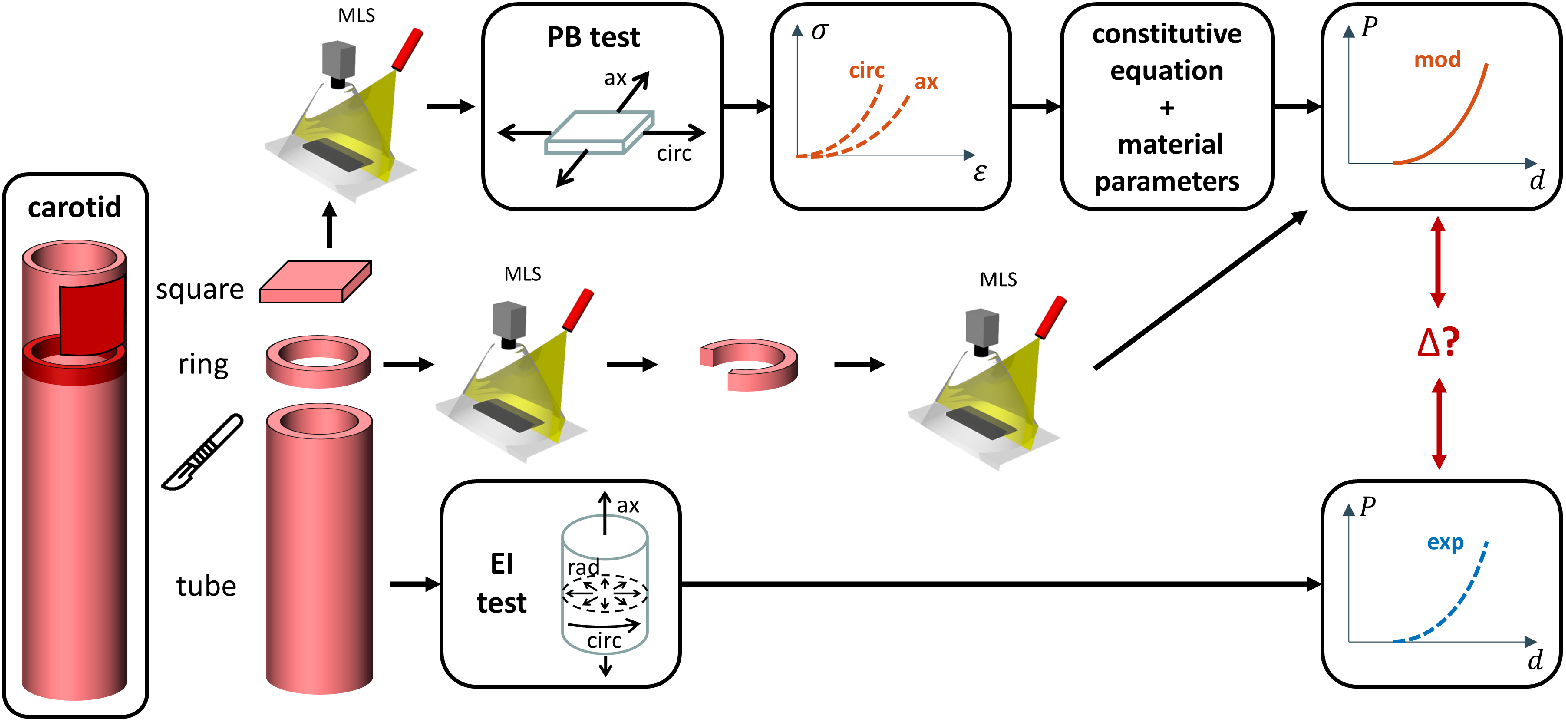
Experimental and Modeled Biomechanical Characterization Workflow. Ovine carotid arteries were prepared for planar-biaxial (PB) loading, ring-opening analysis, and extension-inflation (EI) testing. The axial and circumferential stress–stretch (σ–*λ*) data from PB testing are used to fit material parameters of a chosen constitutive description. These parameters, combined with wall prestretches derived from the ring-opening analysis, enable the modeling (mod) of pressure–diameter (*P*–*d*) behavior. The modeled biomechanical response is then compared to the experimental (exp) *P*–*d* curve obtained from EI testing on the same artery. All geometrical measurements were acquired using micro laser scanning (MLS).

### 2.1. Experimental Workflow

Three complementary types of mechanical tests were performed in this study. Planar-biaxial tests were conducted to yield experimental stress–stretch curves, characterizing tissue behavior under controlled axial and circumferential loading conditions. Ring-opening tests were conducted to assess the opening angle, providing insights into the residual stress state and enabling estimation of *in vivo* arterial prestretches [8]. Extension-inflation tests were performed to generate experimental pressure–diameter data, reflecting the physiological response of the vessels under axial loading and internal pressurization.

#### 2.1.1. Tissue Collection

An overview of the tissue preparation and mounting process is shown in Fig. 2 [9]. Five full-length and fresh ovine carotid arteries, labeled A through E, were collected from one-year-old female Swifter sheep weighing between 50 and 70 kg. The study was approved by the UZ Leuven ethical committee for animal experimentation (ECD) under protocol number 189/2021. The tissues were harvested and immediately preserved in phosphate-buffered saline (PBS) at −80°C until further use [10]. Prior to testing, samples were thawed gradually in the fridge at 4°C. Specimen B is shown as a representative example.

**Figure 2:**
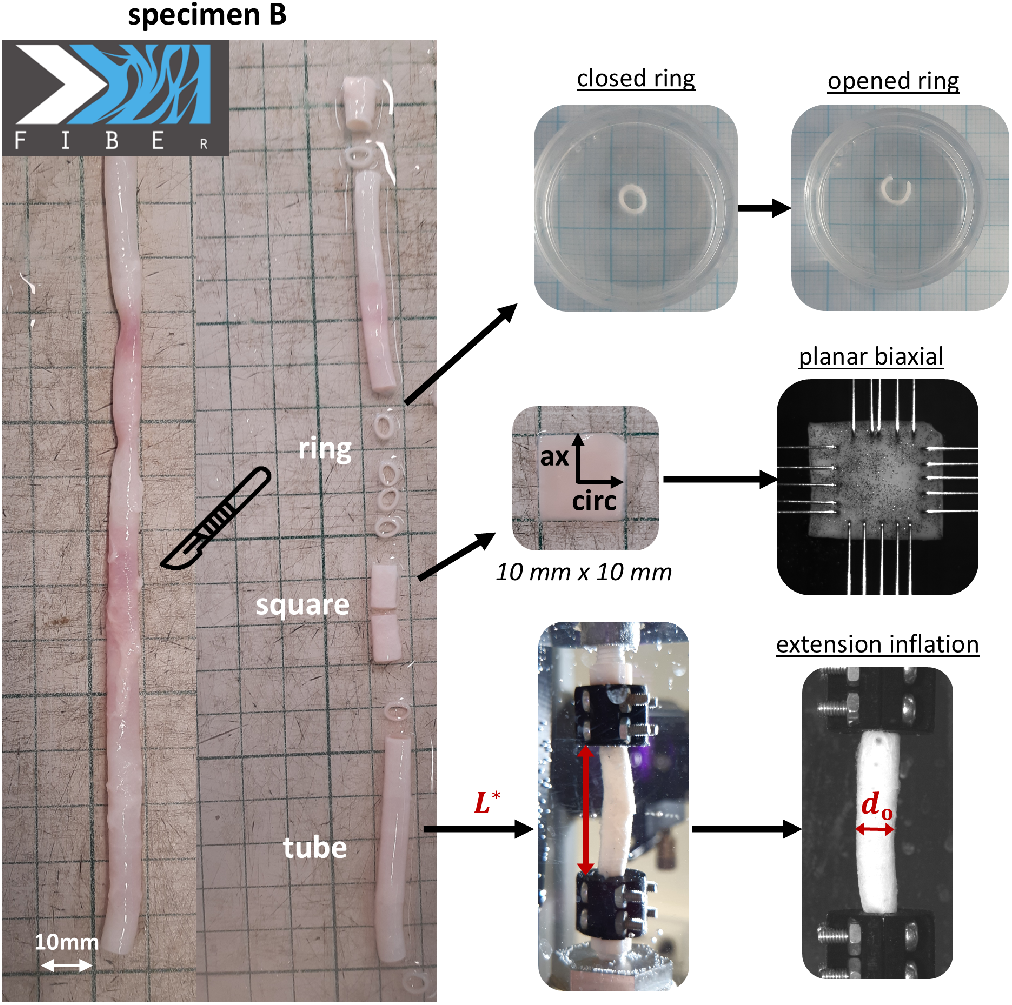
Schematic Overview of Tissue Harvesting and Experimental Mounting. Three test samples were prepared from representative specimen B: a square section for planar-biaxial testing, a ring segment for ring-opening analysis, and a tubular segment for extension-inflation testing. The planar-biaxial configuration includes both unmounted and mounted states, with rakes applied in the axial and circumferential directions to impose controlled deformations. The extension-inflation setup illustrates the mounted length *L*\* and the resulting pressurized outer diameter *d*_o_ during testing.

#### 2.1.2. Planar-Biaxial Testing: Experimental Stress–Stretch

We performed planar-biaxial tensile tests using a ZwickRoell biaxial testing machine for biomaterials on square samples (Table 1). Five rakes were mounted on each side of the sample, equally spaced, resulting in an initial distance of 7 mm between opposite rake sets in both the axial and circumferential directions. The samples were submerged and tested in a 0.9% saline solution, heated to 37°C. During tensile loading, 30% and 50% stretch levels were applied to the rake holders, calculated relative to their initial positions. Stretches were increased stepwise at a rate of 5%/s until single-rake failure occurred. The planar biaxial setup is equipped with four load cells, each with a capacity of 200 N. The load cells have a resolution of 0.0001 N, and a class 1 accuracy is ensured by the calibration certificate until 0.2% of the capacity of the load cells (0.4 N). This means that the lowest readouts of the load cells have a relative deviation and repeatability precision of 0.004 N (1%), a relative resolution of 0.002 N (0.5%) and a relative zero deviation of 0.0004 N (0.1%). Classification 1 remains valid for loads above the accuracy threshold.

**Table 1:** Material Parameters for Square, Ring, and Tube Configuration. We show the data for representative sample B, including the unloaded experimental dimensions for the three geometries (square, ring, and tube), the fitted GOH material parameters, and the calculated prestretches obtained from the ring-opening experiments.

| parameter | definition | value |
| --- | --- | --- |
| <u>square → planar-biaxial</u> |  |  |
| $W_1$ | average sample width, axial direction | 9.9 mm |
| $W_2$ | average sample width, circumferential direction | 10.8 mm |
| $T$ | average sample thickness | 0.30 mm |
| $f_{\text{pre}}$ | tare load | .007 N |
| $c_1$ | neo-Hookean parameter | 49.2 kPa |
| $k_1$ | fiber stiffness coefficient | 5.3 kPa |
| $k_2$ | fiber stiffening coefficient | 9.318 |
| $\kappa$ | fiber dispersion | 0.078 |
| $\alpha$ | fiber angle (w.r.t. the circumferential direction) | 0.0° |
| <u>ring → ring-opening</u> |  |  |
| $R_i^{\text{open}}$ | inner radius, open ring | 2.40 mm |
| $R_o^{\text{open}}$ | outer radius, open ring | 3.15 mm |
| $R_i^{\text{clos}}$ | inner radius, closed ring | 2.45 mm |
| $R_o^{\text{clos}}$ | outer radius, closed ring | 1.83 mm |
| $\beta$ | opening angle | 77° |
| $g_{\text{circ},i}$ | circumferential prestretch, inner wall | 0.97 |
| $g_{\text{circ},o}$ | circumferential prestretch, outer wall | 0.99 |
| $g_{\text{ax}}$ | axial prestretch | 1.04 |
| $D_o$ | load-free outer diameter | 4.90 mm |
| $H$ | wall thickness | 0.62 mm |
| <u>tube → extension-inflation</u> |  |  |
| $L$ | initial tube length | 52 mm |
| $L^*$ | mounted length, measured | 26.49 mm |
| $l^{P=0}$ | unpressurized length, axially loaded | 46.14 mm |
| $D_o$ | mounted load-free outer diameter | 5.49 mm |
| $d_o^{P=0}$ | unpressurized outer diameter, axially loaded | 4.99 mm |
| $\lambda_{\text{ax}}$ | applied axial prestretch | 1.75 |

For each loading level, we applied three axial-to-circumferential stretch ratios {1:1, 2:1, 1:2} using displacement-driven actuators [11]. A 1:2 ratio indicates that half of the 30% or 50% stretch was applied in the axial direction, while the full stretch level was applied in the circumferential direction. Four preconditioning cycles were performed for each loading step and stretch ratio, and the fifth cycle was used for post-processing [12], with a machine sampling rate of 20 Hz.

Experimentally measured forces were converted to engineering stresses using the initial load-free thickness and areal sample measurements, obtained prior to testing via micro laser scanning (Acacia Technology and Gocator Emulator, LMI Technologies). The results from opposite actuators were averaged to obtain single axial and circumferential stress–stretch curves. Instead of considering stretches and displacements at the level of the rakes, we determined the actual tissue stretches by planar particle tracking of the graphite-powder speckle pattern applied to the sample. Digital image correlation (DIC) was performed using the VIC 2D software (Correlated Solutions, isi-sys) by analyzing the speckle pattern at a 20 Hz camera (Manta G-917, Allied Vision) recording rate [13]. The center of the camera was aligned with the zero position of the actuators, imaging orthogonal to the middle of the sample. Note that we consistently speckled and captured the intimal side of the artery. To ensure uniform sample strains, only the central 25% of the loaded area was considered [12, 14].

The zero-strain state for each planar-biaxial stress–strain curve is defined as the point where both axial and circumferential stretches are unity (*λ*_ax_ = *λ*_circ_ = 1), and can be identified during post-processing for each loading ratio by analyzing the last 50% loading curves. Specifically, we chose the planar-biaxial zero-strain state to correspond to the moment when the measured forces exceed a sample-specific predefined threshold. This tare load *f*_pre_ is determined at the start of the experiment by averaging the force values from the four load cells immediately after rake insertion. The mounted configuration is therefore assumed to be the best approximation of a flat, undeformed sample. As our protocol is displacement-driven, preconditioning will consistently lead to negative forces when the actuators move to their initial position, and subsequently exceed the average mounting force or preload *f*_pre_. Since the measured forces in the axial and circumferential direction are not independent, they also share the same tare force to determine a unique zero-strain state for each loading ratio or test cycle. Note that the choice of the tare load or zero-strain state is completely decoupled from the testing protocol, in order to prevent any preliminary assumptions during the planar-biaxial loading itself.

#### 2.1.3. Opening Angle Measurements

Due to the release of residual stresses during planar-biaxial tissue specimen cutting, the reference state is different from the *in vivo* state. We performed opening angle tests on estimated 1 mm thick ring cuts to determine any circumferential prestretch in the arterial wall. The intact rings were submerged in 0.9% saline and scanned in a micro laser scanner (Acacia Technology). The rings were then cut axially, submerged to reduce frictional forces, and ensuring complete residual stress release. A second scan then allowed us to assess the sample-specific ring opening angle. All measurements were performed in the Gocator Emulator software (LMI Technologies).

The closed ring often had an elliptical shape instead of a perfect circle. The major and minor axes of a rectangular region box were then manually adapted to match the inner side of the ring, and averaged to determine a single value for the inner diameter 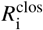, as shown in Fig. 3. The outer diameter of the closed ring 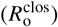 is measured using the same method. For the open ring, inner and outer diameters 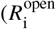 and 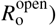 are measured using a circular sector-shaped bounding box. The sector region is delimited by lines extending from the center to the edges of the arc, with the opening angle *β* corresponding to the excluded arc angle. The inner diameter is the largest possible diameter within the open ring, and the outer diameter was taken at the same location.

**Figure 3:**
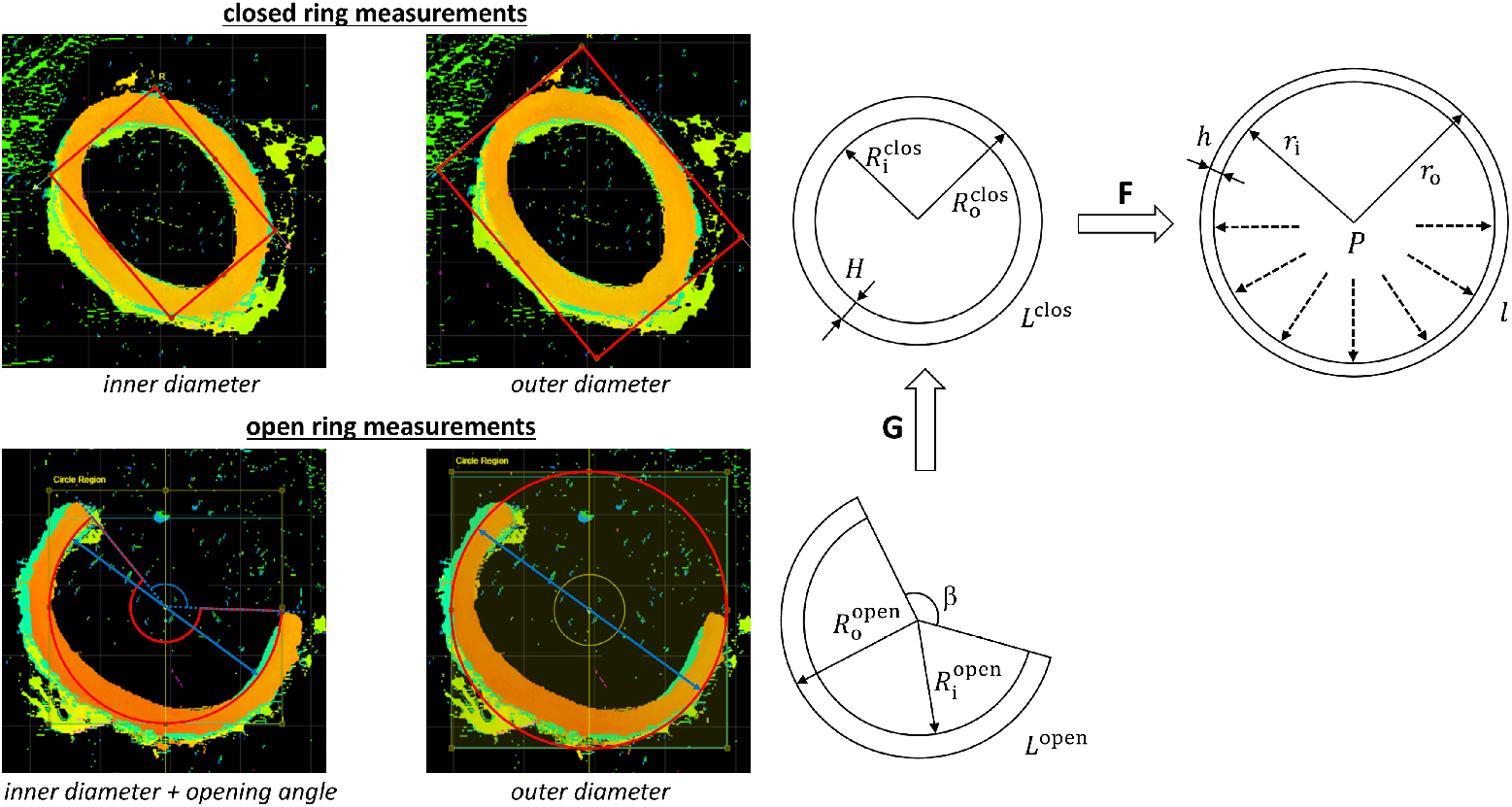
Ring-Opening Experiments and Sample Deformations. Experimental methodology used to assess residual stresses and wall prestretches in tubular arterial tissue. The closed ring configuration is analyzed by fitting a rectangular bounding box to the tissue cross-section, from which the inner and outer diameters 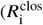 and 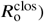 are computed, and the load-free wall thickness *H* can be derived. In the open ring configuration, *β* defines the opening angle. The inner diameter 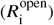 is computed as the largest inscribed circle within the open ring, and the outer diameter 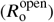 is measured at the same radial location.The prestretch tensor **G** links the reference configuration of length *l*^clos^ to a deformed configuration via the deformation gradient **F**. Axial stretch leads to a deformed length *l* and thickness *h*, with an inner pressure *P* inducing deformed diameters *r*_i_ and *r*_o_.

A flattened square-shaped sample obtained from a curved tubular geometry theoretically introduces a flattening stretch and stress in the circumferential direction, unless the opening angle is 180°, which is assumed to be highly unlikely. Note that the sample is still unloaded in the axial direction, and axial stretch and stress thus remain unchanged. For calcified arteries with a small opening angle, flattening could be substantial. However, in the considered carotid artery samples, the excised square samples flattened completely on their own when laid down freely. Moreover, flattening has been shown to minimally affect the inferred stress–strain pairs [15]. We therefore do not account for flattening effects when correlating the tubular geometry to planar-biaxial tissue testing in our workflow.

#### 2.1.4. Extension-Inflation Tests: Experimental Pressure–Diameter

The experimental extension-inflation tests were conducted using the ZwickRoell testing machine for *tubular* biomaterials. The device is controlled through a custom test environment created within the TestXpertII software. Two actuators of the machine are employed: (I) the linear displacement mechanism, oriented vertically, used for extending the sample and measuring the axial reaction force via a load cell, and (II) the pump, which is utilized to inflate the tubular sample, by pumping water into it, while simultaneously measuring the applied pressure.

To mount the sample, the top end is first fixed between a mounting pin and two mounting clamps, secured with four screws to ensure an airtight seal. The clamps have an axial length of 12 mm and an inner diameter of 4–6 mm, depending on the considered tube dimensions. The assembly lowered until the distance between the underside of the top clamps and the stud of bottom mounting pin equals the initial axial hanging length of the tube sample. The bottom end of the sample is then pulled over the bottom pin. The bottom of the tube is then clamped with similar mounting pieces and screws. The whole sample is submerged in a 0.9% saline solution by raising a 37°C heated bath, and the axial load cell is zeroed. In the absence of axial forces, the measured distance between the clamps in this configuration is equal to the load-free sample length *L*^*^ (see Fig. 2).

Three consecutive load steps were performed, each applying a different axial stretch corresponding to 30%, 50%, and 75% of the initial mounted length. The mounted length is measured with a caliper before testing, and is always smaller than the initial tube length *L* because of the 12 mm clamp dimension on each side. The ultimate axial stretch level of 75% represents physiological axial loading conditions for carotid arteries [16]. An internal pressure of 90 mmHg was applied during the first two axial stretch steps, while the physiological loading step was conducted at 180 mmHg. The latter reflects the upper limit of systolic blood pressure regimes [17]. Lower pressures were used in the initial steps to mitigate the risk of premature tube sealing, loosening, or rupture. Axial loading was applied at a rate of 2.5%/s, and internal pressure was increased at a rate of around 40.0 mmHg/s, with a pump speed of 3.5 nl/min volume. The relative pressure between the inside and outside of the sample is measured directly by the pressure sensor of the test device, with a precision 0.2% of the nominal value. In the extension-inflation test setup, we used an axial load cell of 20 N with a resolution of 67 µN.

The experimental outer diameter displacements are captured using a Grasshopper3 camera (Teledyne FLIR). The imaging device is mounted horizontally and aligned with the center of the sample, with respect to both its length and its circumference. The camera, controlled by the VIC Snap software, was configured at a recording rate of 20 Hz (Correlated Solutions, isi sys).

The outer diameter of the sample in the deformed configuration (*d*_o_) is obtained by processing images captured by the camera in Matlab (The MathWorks, Inc.). We differentiate the mounted, load-free outer diameter *D*_o_, from the unpressurized but axially stretch outer diameter 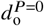, as highlighted for the measured tube in Table 1. We semi-automatically segmented each image to extract the sample and calculate the sample height and width in pixels. We convert pixels to millimeters using the pixel resolution, derived by dividing the known length or height of the mounted sample by the number of pixels in the sample length at the first image of the test protocol.

The converted width or outer diameter is recorded at various points along the sample length. To avoid boundary effects, diameter values in the middle of the sample are reported here. For the experimental pressure–diameter curves, we select the data from the 75% axial stretch loading condition. This step comprises five pressurization cycles, where the first four serve as preconditioning of the sample and minimize viscoelastic effects. Only the stretch or inflation part of the final cycle is used for analysis.

### 2.2. Modeling Workflow

#### 2.2.1. Kinematics & Assumptions

In what follows, we will use uppercase to represent initial reference configurations through Lagrangian variables *X*. Current or deformed configurations will be represented by lowercase variables *x*, as will model parameters. Tensors are denoted in bold upright font, vectors in bold italic, and scalars in italic. Subscripts and superscripts representing indices are italicized, while other subscripts and superscripts, such as abbreviations, are set in upright font.

We define the deformation gradient as **F** = ∂*x* /∂ *X*, where tissue incompressibility implies that the Jacobian *J* = det (**F)** = 1 [18, 19]. We will highlight planar-biaxial configurations through subscript PB, while we will refer to extension-inflation setups as EI, resulting in pressure–diameter results (*P*–*d*). For consistency, we will write all modeled deformations as **F**^mod^, and the corresponding modeled First Piola-Kirchhoff stresses as **P**^mod^. Experimentally measured deformations and stresses will be denoted as **F**^exp^ and **P**^exp^, respectively.

In both the planar-biaxial test and the extension-inflation test we ignore shear deformations. In the planar-biaxial test we assume that the artery is consistently stretched in two perfectly orthogonal directions, aligned with the circumferential and axial direction of the artery. Using the stretch definition *λ* = *dx /dX*, we can write **F**_PB_ = diag {*λ*_circ_, *λ*_ax_, *λ*_rad_}, in a Cartesian coordinate system, whereby the three principal directions align with the axial, circumferential and radial direction of the artery, respectively. In the extension-inflation test, we assume an axisymmetric radial pressurization accompanied by an axial elongation, yielding a deformation gradient **F**_EI_ = diag{*λ*_rad_, *λ*_circ_, *λ*_ax_}in a cylindrical coordinate system. In both tests, the radial stretch is imposed by incompressibility, 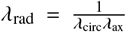 whereas the circumferential and axial stretches will be imposed by the testing device.

Note that we disambiguate stretch from strain by defining the nominal or engineering strain as *e* = (*dx* −*dX*)/*dX* = *λ*−1.

#### 2.2.2. Constitutive model

Assuming hyperelastic material behavior, the Cauchy stress may be written as a function of the deformation as

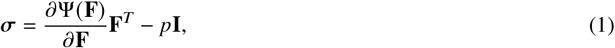

where Ψ is the strain energy density function, and *p* is a Lagrange multiplier enforcing the incompressibility constraint, which will be determined by specific boundary condition assumptions of the planar-biaxial or extension-inflation setup. Subsequently, we can calculate the corresponding first Piola-Kirchhoff stress (i.e. the transpose of the engineering stress) as **P** = *Jσ***F**^−*T*^.

We adopt the Gasser–Ogden–Holzapfel (GOH) strain energy density function, which assumes Fung-type strain-stiffening fiber families embedded in an isotropic matrix [20–23]. From the right Cauchy-Green deformation tensor (**C** = **F**^*T*^ **F**), we can define the first and fourth strain invariants used in the GOH constitutive equation as 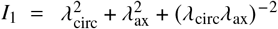 and 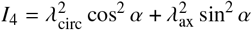, respectively. *λ*_ax_ and *λ*_circ_ are again the principal stretches in the axial and circumferential direction. The angle *α* is defined as the collagen fiber orientation with respect to the circumferential direction in the circumferential–axial plane, assuming two symmetrically oriented fiber families. Due to this symmetry, the fiber angle *α* is restricted to 0°≤ *α* ≤90°. As such, the GOH model combines an isotropic neo-Hookean contribution based on the first invariant with an anisotropic fiber contribution of the fourth invariant, modeled by a quadratic-exponential function.

The resulting strain energy density function of the GOH constitutive model can be written as

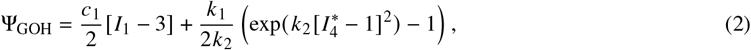

with the shear modulus *c*_1_ *>* 0, the stiffness-like parameter *k*_1_ *>* 0 and the non-dimensional coefficient *k*_2_ *>* 0. The latter modulates the exponential strain-stiffening characteristics of the collagen fibers. The mixed isotropic-anisotropic fiber invariant term expands as 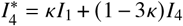, where the microstructural fiber dispersion parameter *κ* modulates the degree of isotropy of the collagen fibers. When *κ* = 0, we have 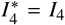, and the model reduces to non-dispersed collagen fibers along the direction defined by the angle *α*. When *κ* = 1 /3, we have 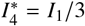, and the model reduces to a fully isotropic model.

#### 2.2.3. Material Parameter Optimization

To identify the five optimal material parameters {*c*_1_, *k*_1_, *k*_2_, *κ, α*} of the GOH model, we perform a parameter optimization using the lsqnonlin routine in Matlab [24, 25]. The optimization minimizes the 2-norm of the residuals between experimental and model-predicted stress components:

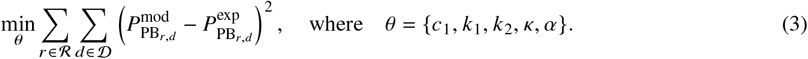

The optimization combines the three loading ratios, ℛ= {1:1, 2:1, 1:2}, in the axial and circumferential direction,𝒟= {ax, circ}, of the fifth loading cycle at 50% stretch. To balance the contributions of each experimental direction and loading ratio, we applied a scaling to normalize their magnitudes. Per sample, we weighted the error during fitting per ratio and per direction, in order to account for different data ranges equally. Moreover, we also compensated for the number of experimental data points per curve. Convergence to local minima was avoided by means of ten different initial guesses in a multi-start strategy. The Lagrange multiplier *p* in 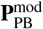 was calculated through the assumption of zero radial stresses 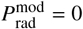.

#### 2.2.4. Simulating Pressure–Diameter Behavior

The analytical framework assumes a thick-walled, axisymmetric, cylindrical artery. The load-free configuration of an extension-inflation, at which **F**_EI_ = **I**, corresponds to the closed configuration at zero pressure, while the deformed state represents any pressurized configuration of the artery [26, 27]. However, arterial tissue in its cylindrical shape is still in a prestretched state. Figure 3 shows the link between the deformation gradient **F**_EI_, and the prestretch tensor**G** = diag {*g*_circ_, *g*_ax_, *(g*_circ_*g*_ax_)^−1^ }, recalling the incompressibility constraint. **G** is used to disambiguate the undeformed load-free configuration from the reference stress-free configuration. From the opening angle *β*, we can compute

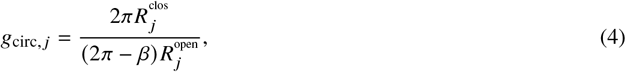

at every integration point *j* through the thickness. A unique value of *g*_ax_ is then iteratively found by matching the experimentally measured load-free thickness 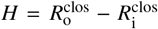 with the modeled radial coordinates through the thickness in the same reference configuration, i.e. for an inner pressure *P* = 0 and an axial stretch *λ*_ax_ = 1.0.

The total deformation of the artery is then equal to 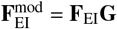, with the prestretch tensor **G** determined in the closed, load-free configuration, and the extension-inflation deformation gradient **F**_EI_ dependent on the applied inner blood pressure and imposed axial stretch. Note that, since we assume a thick-walled cylinder, both tensors also vary along the arterial wall thickness.

Indeed, when applying a constant axial stretch *λ*_ax_ = *l*/*L*^clos^, the remaining unknown in 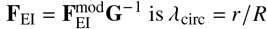, which is dependent on the radial position. *R* _*j*_ and *r* _*j*_ can represent any corresponding undeformed and deformed radial coordinates in the arterial wall thickness. Consequently, we define *λ*_circ, *j*_, with *j* referring to a specific integration point through the wall.

A blood pressure *P* is applied on the inside of the cylinder, with the outside pressure always taken as zero. By considering axisymmetric constraints and ignoring accelerations and body forces in the equation of balance of momentum 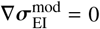 the following equation for stress equilibrium in the radial direction can be derived [28]:

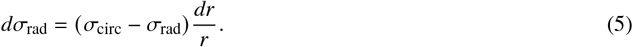

Integration over the wall, i.e. from the inner to the outer radius, and knowing that σ_rad_ (*r*_o_) = 0 and σ_rad_ (*r*_i_) = -*P*, yields

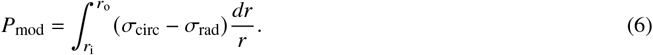

Consequently, the inner circumferential stretch *λ*_circ,i_ for any applied blood pressure *P* can be computed using following objective function:

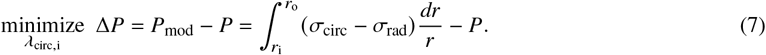

An initial guess of *λ*_circ,i_ is required in the optimization scheme, and should be adjusted to match the expected deformations in order to enhance convergence over the complete pressure range. Volume conservation allows to derive *λ*_circ,j_ = *r* _*j*_/*R* _*j*_ at any point in the wall. Indeed, due to incompressibility, it holds that

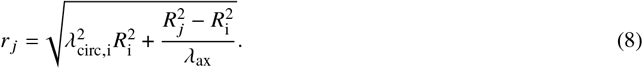

Now, when calculating the modeled extension-inflation Cauchy stresses 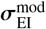 through Eq. (1), the Lagrange multiplier *p* varies throughout the wall too. For the inner wall, it holds that 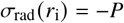. Substitution yields an expression for *p* at the inner wall. Similarly, 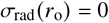. Therefore, to find the solution for the Lagrange multiplier within the wall, we consider that Eq. (6) should hold at every radial position. Thus,

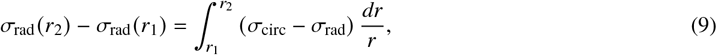

should hold for all radial positions *r*_1_ and *r*_2_ between the inner and the outer arterial wall, *r*_i_ and *r*_o_. This allows us to obtain *p* at every radial position. Here, the integral was solved through 50 integration points and a trapezoidal method with equal spacing. Solving equations 7, 8 and 9 allows us to derive the corresponding 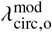 for any given pressure and applied axial stretch *λ*_ax_, from which also the resulting outer diameter *d*_o_ can be obtained. As such, we report the modeled pressure–diameter behavior for each specimen, over the range 0–200 mmHg.

The resulting axial Cauchy stress, averaged through the wall thickness, is equal to

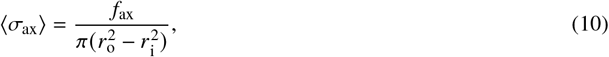

where *f*_ax_ is the axial load on both ends of the tube in order to maintain the deformation [19]. Taking into account the contribution of the pressure on the end area of a tube with closed ends, we can then define the reduced axial force 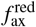 as

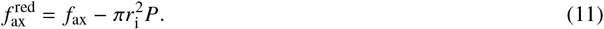

#### 2.2.5. Extension-inflation optimization

From the modeled and experimental outer diameters, an additional optimization scheme can be performed through

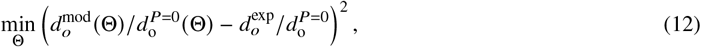

over the entire 0–200 mmHg pressure regime. We minimize the relative outer diameters by normalizing each value with respect to its unpressurized diameter. The pressure–diameter data from 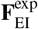 or 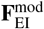is always considered at axial stretch *λ*_ax_, i.e. at deformed length *l*. In the optimization procedure, an either-or approach is adopted, meaning that only one parameter, or subset of parameters, is optimized at a time, while the remaining variables are held fixed at their initial values listed in Table 1. Accordingly, different optimization schemes can be defined as:

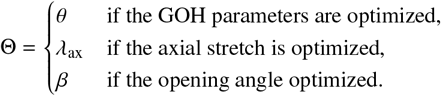

Recall that θ is the set of GOH parameters, θ = {*c*_1_, *k*_1_, *k*_2_, *k, α*}.

## 3. Results and Discussion

In what follows, we will systematically present the results according to our workflow by first reporting the planar-biaxial stress–stretch experimental curves in the axial and circumferential direction. We overlay the experimental results with the simulated planar-biaxial curves computed by the Gasser–Ogden–Holzapfel (GOH) constitutive model fitting. Subsequently, the extracted material parameters are used to simulate the modeled pressure–diameter response of an idealized cylindrical geometry, and to compare the results with experimental extension-inflation data. We then investigate the capacity of the GOH parameters to fit the extension-inflation data directly, and finally assess the sensitivity of the axial stretch *λ*_ax_ and opening angle *β*. Due to the large variability between biological subjects and the sample-dependency inherent in our workflow, statistical analysis was deemed inappropriate. Instead, a single sample was selected as use case and proof-of-concept for our data analysis. The Appendices contains the results for all the remaining samples.

### 3.1. Biaxial Fitting with GOH Material Model

The resulting experimental engineering stress and DIC-derived stretch curves for the three loading ratios, based on the final loading cycle after preconditioning, are presented for representative sample B in Fig. 4. Table 1 reports the undeformed square dimension of the planar-biaxial samples, used to transform measured actuator forces into first Piola-Kirchhoff stresses. According to Section 2.2.2, the material parameters for the GOH constitutive model can then be fitted to calculate the modeled stresses, 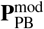, in the corresponding strain regimes. *R*^2^ = 0.908 gives an indication of the goodness of fit between the experimental and modeled stress–stretch curves per sample, for the two directions and three loading ratios combined. The model’s axial and circumferential predictions for the three loading ratios are overlaid with the corresponding experimental curves, demonstrating the model’s ability to capture the anisotropic hyperelastic mechanical behavior of the arterial tissue. From Fig. 4, we note an observably stiffer experimental mechanical behavior in the circumferential direction with respect to the axial direction, consistent with the resulting GOH material parameters. Indeed, both modeled collagen fiber families were found to be oriented predominantly in the circumferential direction, exhibiting a small fiber dispersion. The latter is also in agreement with microstructural fiber analysis in arteries, with a probability density function of collagen orientations towards the circumferential direction, despite a large spread and uncertainty [29].

**Figure 4:**
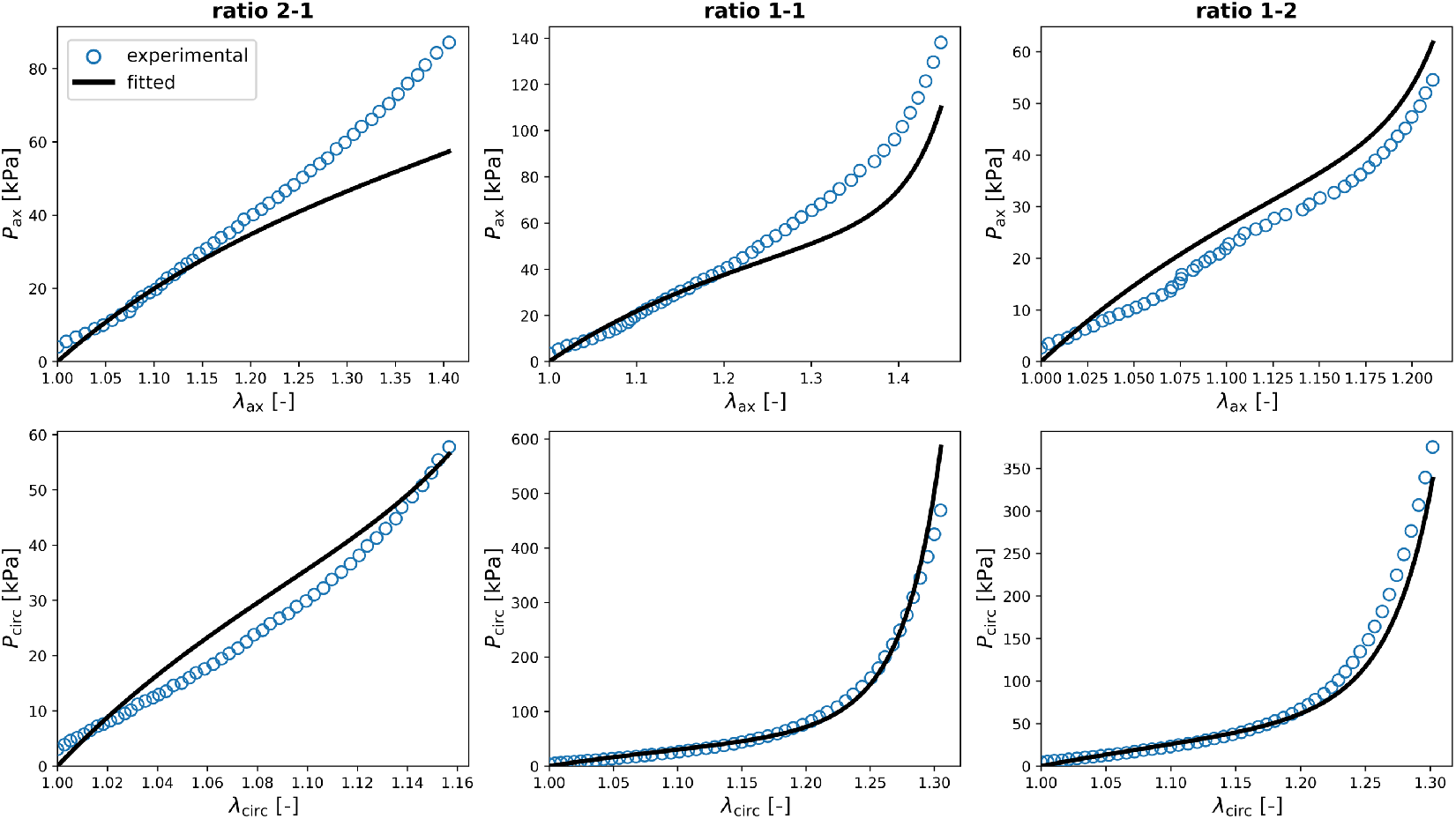
Experimental and Modeled Planar-Biaxial Stress–Stretch Curves. Axial and circumferential first Piola-Kirchhoff stresses (*P*_ax_ and *P*_circ_) plotted for sample B under three loading protocols with stretch ratios of 2:1, 1:1, and 1:2 in the axial and circumferential directions (*λ*_ax_ and *λ*_circ_). Experimental stress–stretch data are shown alongside the modeled stress responses obtained using the fitted material parameters of the GOH constitutive model. The goodness of fit between modeled and experimental biaxial data of the three ratios and two directions is given by *R*^2^ = 0.908.

#### Retrieving high-quality and biofidelic experimental planar-biaxial data is challenging

The observed tissue deformations are considerably lower than the nominal 50% stretch applied at the rakes. This discrepancy arises from two factors: the actual strains at the center of the sample are smaller than those at the rakes [12], and our methodology to define the zero-strain state. Indeed, the initial starting point of the stress–stretch curve is set at the average force recorded by the four load cells immediately after mounting, prior to the onset of preconditioning. This approach provides the best available approximation of a flat, unstressed planar-biaxial tissue configuration. A fixed preload of 0 N should be avoided because of inevitable mounting forces. Defining zero-strain at higher forces would shift the origin of the stress–stretch curve even further along the loading axis, thereby omitting an additional portion of the low-strain response and overestimating tissue extensibility. Note that inelastic deformations can be present in part of the loading protocol, both during preconditioning and the main loading sequence. This is further evidenced by negative force values observed when the rakes return to their initial positions. Occasional discontinuities in the experimental stress–stretch curves are present, particularly in regions where the DIC strain measurements appeared challenging due to lighting, speckle tracking errors, or local sample deformations. The latter is also highlighted by a small but acceptable amount (*<*10%) of undefined values in the DIC-derived stretches. These limitations underscore the difficulty in obtaining clean, artifact-free biaxial data from soft biological tissues.

#### The fiber-dispersed GOH material model fits planar-biaxial arterial data well

Through the material parameter optimization of Eq. (3), the fiber-reinforced GOH model achieves an overall fitting performance of *R*^2^ *>* 0.8 for all samples, indicating a consistent good fit between experimental data and the model stress–stretch predictions. Notably, the isotropic matrix stiffness parameter *c*_1_ appears to scale with the exponential strain-stiffening coefficient *k*_2_ of the collagen fibers, while the fiber stiffness parameter *k*_1_ exhibits an inverse trend. This suggests a compensatory behavior between the isotropic and anisotropic components of the model. The fitted fiber direction of both collagen families is entirely oriented along the circumferential direction (*α* = 0°), but an axial stiffness contribution is introduced through a non-zero dispersion parameter (*κ >* 0). While this captures physiologically plausible mechanical behavior, it can also reveal over-parametrization of the GOH model in certain cases [23]. Actually, the structural parameters *α* and *κ* could be derived from microstructural analysis, such as second harmonic imaging through multi-photon microscopy. However, literature has shown that it remains challenging to accurately determine collagen fiber angles and dispersions in arterial tissue [30]. Alternatively to the conventional GOH model, constitutive artificial neural networks can be implemented to autonomously learn material behavior from the biaxial stress–stretch data [31–34]. This data-driven framework bypasses predefined model assumptions, and provides additional insights into stress–stretch behavior, offering a powerful alternative for capturing complex arterial mechanics. Instead of relying on the GOH strain energy density function, we explored automated model discovery in the Appendices (Fig. A8), and reported an updated constitutive description for carotid arteries.

### 3.2. Comparing Pressure–Diameter Responses

In this section, we analyze the pressure–diameter relation, Fig. 5a, as well as the relative circumferential stretch, Fig. 5b. The circumferential stretch in the extension-inflation plot is defined as the relative outer diameter, and is computed as 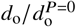. Note that both the current and initial diameter are computed in a deformed axial stretch state, i.e. with an applied *λ*_ax_. Table 1 reports the experimental length and outer diameter of the tube, together with the ring measurements used to retrieve the model’s load-free diameters and corresponding wall thickness at zero pressure. The table also shows the measured opening angles and corresponding circumferential prestretch at the inner and outer wall of the rings, together with their corresponding axial prestretch *g*_ax_. Unless otherwise stated, the absolute and relative modeled curves are always calculated and plotted with an applied *λ*_ax_ = 1.75.

**Figure 5:**
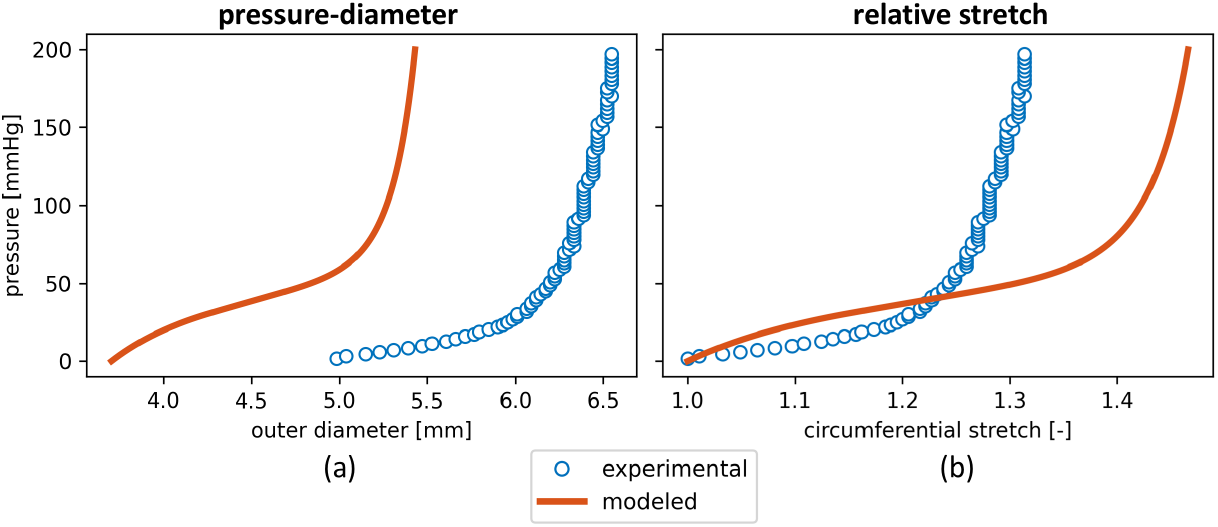
Modeled and Experimental Pressure–Diameter Behavior. Comparison of modeled and experimental pressure–diameter behavior for representative sample B, showing (a) the absolute outer diameter, and (b) the relative circumferential stretch. The modeled curves are generated using GOH material parameters fitted from the corresponding biaxial stress–stretch data. Experimental pressure–diameter data is extracted from the extension-inflation test.

#### The modeled curves underestimate the absolute pressure–diameter behavior

Fig. 5a highlights how the model underestimates the absolute diameter values for almost all samples, except sample E. To get a sense of biofidelity of our results, the *P*–*d* curves could potentially be validated using *in vivo* imaging data, providing diameter measurements at diastolic and systolic pressures. However, such a validation protocol is challenging, as typically only inner diameters can be extracted from standard imaging data, and arterial blood pressures are often not recorded simultaneously with the imaging modality. Intravascular ultrasound has been used to measure both luminal diameters and pressurized arterial wall thicknesses, and may be further explored and implemented [35, 36]. Pre-operative imaging of the sheep was not performed in this study, but the numerical values of the modeled absolute diameters are in agreement with literature data reported for ovine carotid arteries [37]. More specifically, inner diameters between 5 mm and 6 mm have been reported in the literature for systolic and diastolic blood pressures, which aligns well with our results. Normal diastolic and systolic arterial blood pressure values in sheep range from 60–80 mmHg and 90–120 mmHg, respectively [17]. This corresponds to the highly non-linear exponential regime in our *P*–*d* curves, leading to only minor diameter changes and thus a limited pulsatile effect.

#### The modeled curves overestimate the relative pressure–circumferential stretch behavior

Fig. 5a also shows that the zero-pressure diameter does not coincide: the axially stretched but pressure-free outer diameter in the model is consistently lower than the experimentally measured outer diameter, even after an axial stretch of *λ*_ax_ = 1.75. The model’s initial outer diameter and load-free wall thickness are a potential cause of this mismatch, as the closed-ring configuration is a direct input of the simulations, without taking the tube dimensions into account. To overcome this inherent bias, we also assess the pressure–diameter behavior in a relative manner through the circumferential stretch in Fig. 5b. The latter was obtained by normalizing the diameter profile with its unpressurized but axially loaded value. Here we see that the model generally overestimates the relative diameter change compared to the experimental extension-inflation data. The best relative agreement between model and experiment is observed for samples C and D, see Appendix Figs. A1 and A2.

#### Probing relevant physiological stretches remains challenging

Ideally, we aim to reproduce the physiological stretch regime of arteries during biaxial testing, as this will enhance the biofidelity of the mechanical characterization and any downstream modeling or simulation steps. From in-house experiments at the Experimental Cardiac Surgery lab at KU Leuven on similar ovine carotids, we measured an *in vivo* axial length of approximately 3.5 cm reducing to an *ex vivo* length of around 2 cm, suggesting an axial stretch (*λ*_ax_) of about 1.75. Furthermore, *in vivo* echo-ultrasound imaging of a carotid artery under pulsatile pressure showed a loaded inner diameter ranging between 4.5 and 5.5 mm, and a load-free inner diameter between 3.5 and 4 mm, corresponding to estimated circumferential stretches between 1.3 and 1.4. However, achieving such high deformation ranges is limited by experimental constraints, especially the failure of the rakes or clamping system at large displacements. Figure 6 shows how the arterial tissue was loaded in a very different axial-to-circumferential stretch domain when comparing planar-biaxial results with extension-inflation data. The experimental stretches show how 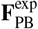 and 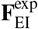 differ, and none of the biaxial stretch ratios come close to the extension-inflation deformations. The axial and circumferential components of the prestretch tensor in **F**_EI_**G** determine the reference configuration of the extension-inflation data (*λ*_ax_ = *λ*_circ_ = 1). Figure 6 also nicely shows the absence of any relative axial stretch during pressurization. Indeed, experimentally, the clamps prevent axial movement and theoretically, the model predicts a constant axial deformation of *λ*_ax_*g*_ax_ ≈ 1.82.

**Figure 6:**
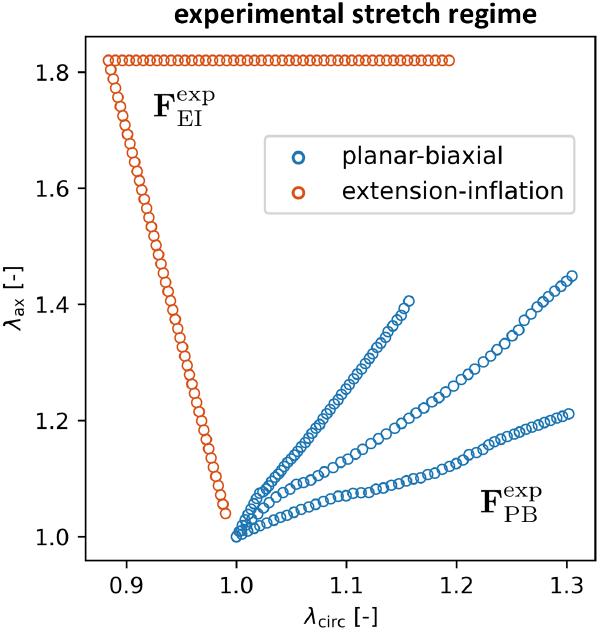
Experimental Biaxial Stretch Regimes. Comparison of the three planar-biaxial axial-to-circumferential stretch ratios 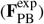 with the extension-inflation experimental deformations.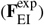 To guarantee an equivalent reference configuration, the stretch domain of the extension-inflation test includes the prestretch tensor **G**, derived from the corresponding opening angle experiments.

#### Model predictions are influenced by both the planar-biaxial dataset and the constitutive description

Equation (3) considers the three axial-to-circumferential stretch ratios from the planar-biaxial dataset. To test if only a selected loading ratio would be more appropriate for accurately representing the *in vivo* loading conditions of a carotid artery, Figs. A3–A5 in the Appendices assess the GOH material fitting and model predictions solely based on one of the 2:1, 1:1 or 1:2 planar-biaxial ratio. Based on the experimental measurements mentioned above, the axial-to-circumferential stretch ratio 2:1 approximates the best the biaxial stretch regime that the sample experiences during extension-inflation. However, this omits the strain-stiffening mechanical behavior of the collagen fibers in the circumferential direction from the planar-biaxial material fitting. When performing material parameter fitting based on planar-biaxial *λ*_ax_ 1.4 → and *λ*_circ_ → 1.2, the pressure–diameter model predictions therefore don’t accommodate the exponential mechanical signature of an extension–inflation test (see Appendix Fig.A3). Planar-biaxial ratios 1:1 and 1:2 both reach *λ*_circ_ → 1.4 and result in a non-linear mechanical signature of the stress–stretch curve, but do not represent a physiological loading condition. Extrapolation of the fitted GOH model and parameters contributes to the poor pressure–diameter match when predicting extension-inflation data, as the simulation uses constitutive parameters fitted on mechanically non-physiological planar-biaxial tensile tests. Rake failure at increased stresses prevented us from extending the stretch regime to *λ*_ax_ → 1.75 and *λ*_circ_ → 1.375, but future planar-biaxial testing protocols and loading ratios should be designed to experimentally reach those tissue-specific *in vivo* loading conditions [38].

In what follows, we will discuss possible solutions to the observed pressure–diameter offsets, by refitting the GOH parameters to the extension-inflation data directly, and analyzing the sensitivity of the axial stretch and circumferential prestretch.

### 3.3. Parameter Optimizations

This section will explore how different modeling strategies and parameter choices can influence the accuracy of simulated pressure–diameter behavior for the representative specimen B. To do so, we performed a least squares optimization of the modeled outer diameters over the whole pressure regime, see Eq. (12). Figure 7 compares several approaches and results. In particular, we will look into an alternative GOH parameter optimization, fitted to the extension-inflation data directly. We will then analyze the sensitivity of the modeled axial stretch and implemented ring opening angle, which influences the circumferential prestretch of the tissue.

**Figure 7:**
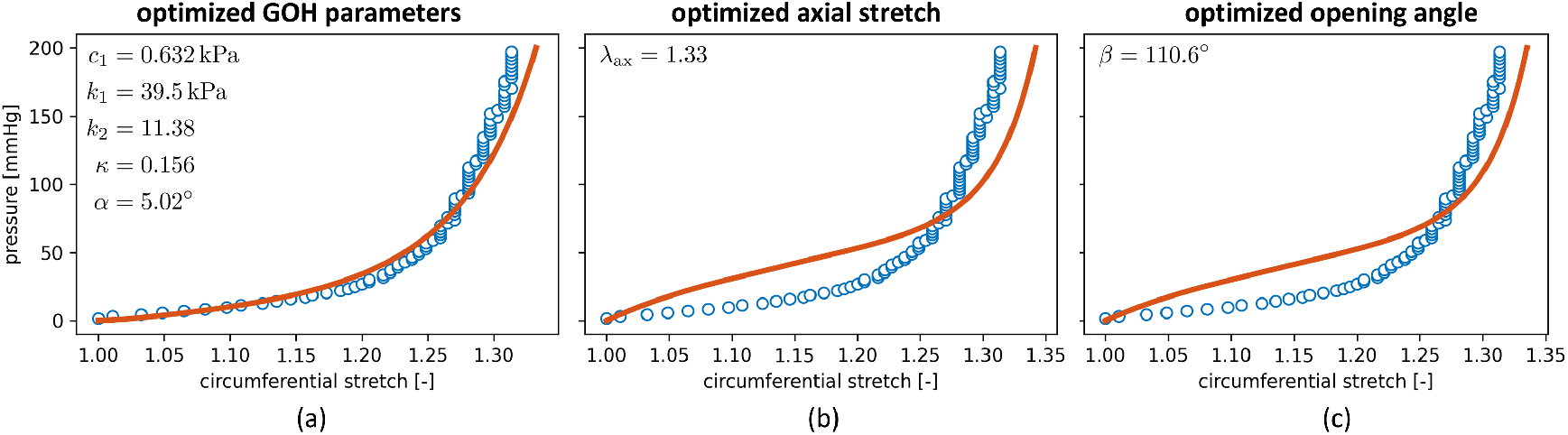
Parameter Optimizations. For the representative case of specimen B, we compare different approaches to match experimental pressure– circumferential stretch behavior. (a) GOH model fit directly on the extension-inflation data. (b) Optimization of the axial stretch (c) Optimization of the circumferential prestretch through the opening angle.

#### 3.3.1. Constitutive Parameter Fitting Based on Extension-Inflation

Because the GOH parameters are fitted to planar-biaxial test data which loads the considered square samples in a very different strain regime compared to the extension-inflation deformations, this leads to non-biofidelic extrapolation of the results. Therefore, we here fit the GOH model directly onto the pressure–diameter data, through the optimization in Eq. (12), for the set of GOH parameters θ = {*c*_1_, *k*_1_, *k*_2_, *κ, α*}, and with *λ*_ax_ = 1.75.

##### The GOH model can closely match experimental pressure–diameter relations as well as planar-biaxial data, but not simultaneously

Directly fitting the GOH model to the experimental extension-inflation data (Fig. 7a) greatly improves the agreement between the relative outer diameter of the model and the experiment. The changes in predicted pressure–diameter behavior originate from notably different fitted material parameters. The shear modulus *c*_1_ decreases drastically, while the fiber stiffness parameter *k*_1_ increases by nearly two orders of magnitude. Additionally, the exponential parameter *k*_2_ increases sharply from 0.0932 to 11.38, indicating a much stronger strain-stiffening effect in the fiber direction. These parameter shifts underscore that planar-biaxial and extension-inflation tests probe fundamentally different aspects of the tissue’s mechanical response. This is also in line with previous research relating *ex vivo* triaxial shear testing to *in vivo* pressure–volume loop data [39, 40]. Nevertheless, both planar-biaxial and extension-inflation approaches here provide valuable insights into the behavior and influence of the GOH parameters under distinct loading scenarios. We could also re-evaluate the modeled planar-biaxial stress–stretch curves with the updated optimized GOH parameters from the extension-inflation data. Appendix Fig. A6 shows an underestimation of the stress in the low-stretch regime, but an overshoot of modeled stress in the biaxial high-stretch regime. This overestimation is likely related to the collagen fiber stiffness and stiffening non-linearity in the updated GOH material properties, dispersed around the circumferential direction, and required to meet the exponential pressure–diameter mechanical behavior.

##### The (an)isotropic balance of the GOH model has an important influence on the convex or concave shape of the pressure–diameter curve

Interestingly, the modeled GOH curve in Fig. 5a exhibits a pronounced concave slope at low pressures (*P <* 50 mmHg), transitioning into a highly non-linear, convex diameter change as pressure increases, characteristic of strain-stiffening behavior. In contrast, the experimental pressure–diameter curve displays a solely convex shape with an elongated toe region of approximately 20% stretch. When reducing the isotropic parameter *c*_1_ of the GOH model while increasing the fiber-related parameters *k*_1_ and *k*_2_, the initial concave behavior of the *P*–*d* curve diminishes, resulting in a fully convex and exponentially increasing mechanical response in circumferential stretch (Fig. 7a). In Eq. (3), the five GOH parameters were scaled to account for relative importance, but a rigorous sensitivity analysis on the pressure–diameter results through Sobel indices could further reveal mutual influence.

#### 3.3.2. Axial Stretch Analysis

The influence of the axial stretch on the model predictions is given in Fig. 8. By varying *λ*_ax_ over a physiologically relevant domain from 1.0 to 2.0, we can observe clear changes in predicted pressure–diameter and circumferential stretch behavior. In Fig. 9, we also introduce the resulting effects on mean axial Cauchy stress and reduced axial force across blood pressures.

**Figure 8:**
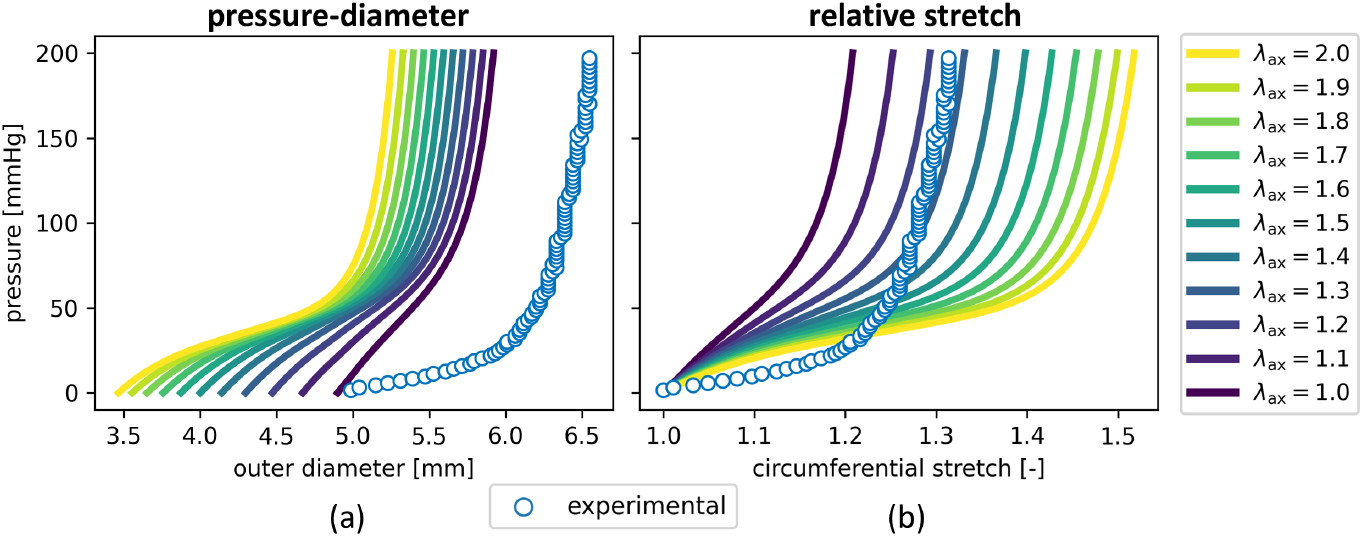
Axial Stretch Sensitivity. Effect of varying axial stretch on model predictions for representative specimen B. Axial stretch values range from 1.0 to 2.0 in increments of 0.1. (a) Modeled pressure–diameter curves for each axial stretch value. (b) Corresponding relative circumferential stretch behavior. The experimentally applied axial stretch during extension-inflation testing was *λ*_ax_ = 1.75, as imposed by the device actuator.

**Figure 9:**
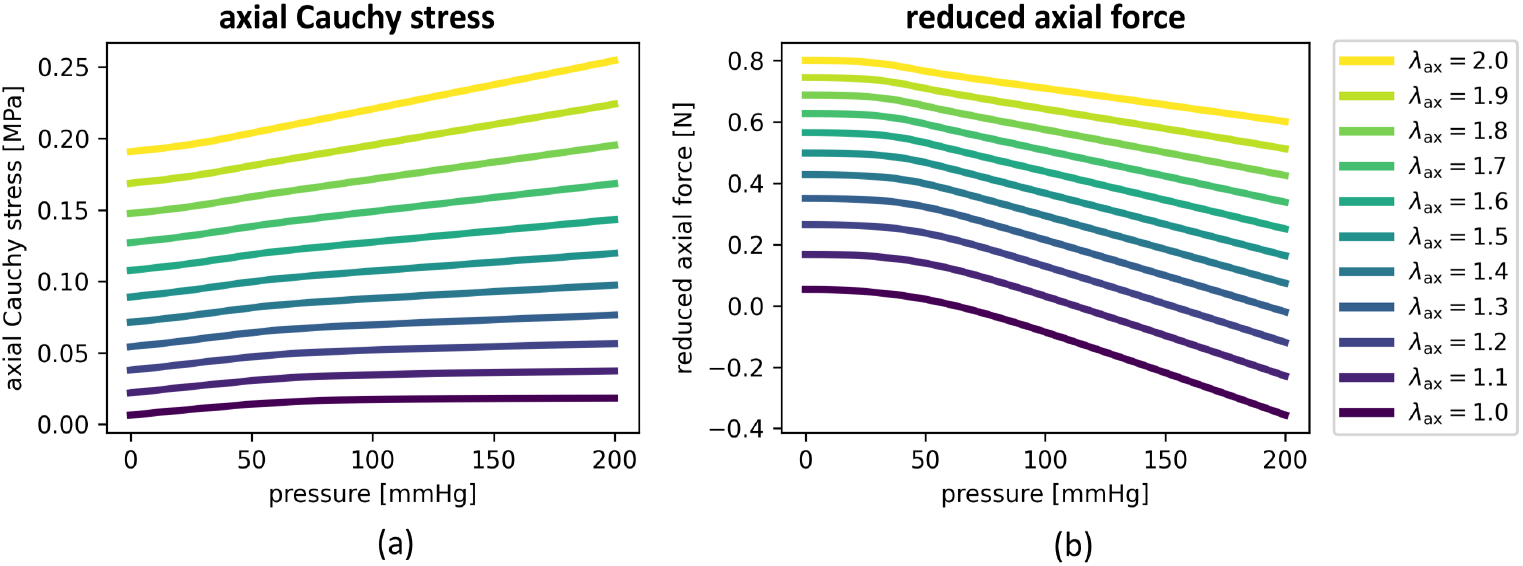
Axial Loading Conditions. Effect of varying axial stretch on resulting stresses and forces for representative specimen B. (a) Resulting axial Cauchy stress ⟨ σ_ax_ ⟩, averaged through the wall thickness. (b) Calculated reduced axial force 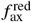. The experimentally applied axial stretch during extension-inflation testing was *λ*_ax_ = 1.75. The optimized axial stretch is *λ*_ax_ = 1.33.

##### Increasing the axial stretch reduces absolute diameters, but increases the circumferential deformation

Interestingly, increasing the modeled axial stretch leads to a decrease in the absolute modeled outer diameter (Fig. 8a), while simultaneously increasing the relative circumferential stretch regime (Fig. 8b). However, increasing axial stretch values to supra-physiological ranges will ultimately result in a severe increase in tissue stiffness and a reversal of the trend. Indeed, *λ*_ax_ *>* 2.5 will cause the relative diameter curve to shift leftwards again with increasing pressures. Ultimately, and in the specific case of sample B here, *λ*_ax_ ≈ 3.75 will match the experimental curve again. However, such an axial stretch range is far above the experimentally observed and reasonable *in vivo* values of *λ*_ax_. A similar relative outer diameter optimization in Eq. (12) can be performed to find the optimal axial stretch. Figure 7b therefore retains the initial GOH parameters from Table 1, but optimizes *λ*_ax_, resulting in a value of *λ*_ax_ = 1.33. The remarkable lower axial stretch improves model performance without re-fitting material constants, highlighting that uncertainties in axial boundary conditions of the tubes can have a strong influence on the model’s accuracy.

##### Axial loading should be included in extension-inflation parameter optimizations

The optimization scheme could be enriched with axial data by normalizing Eq. (12) with the diameters before axial loading, 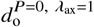. In fact, one should also consider the measured axial forces during the extension-inflation experiment in order to perform a parameter optimization on actual biaxial loading conditions. Appendix Fig. A7 reports the experimentally measured axial forces during the extension-inflation of sample B. Calculated for physiological pressures and at *λ*_ax_ = 1.75, we added the modeled values of *f*_ax_. For the range of varying axial stretches, Figs. 9a and 9b show the modeled Cauchy stress ⟨σ_ax_⟩ and calculated reduced axial load 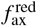, based on Eqs. (10) and (11) respectively. For increasing values of *λ*_ax_, the axial wall stress profile shifts from decreasing to increasing pressure response, and is inversely proportional to the reduced axial force. The axial stretch analysis demonstrates how sensitive wall stress estimation is to longitudinal loading, and highlights the importance of the experimental tube assumptions in the pressure–diameter predictions. In the literature, the most physiological axial stretch is often defined as the value at which the axial load remains nearly constant, despite an increase in intimal blood pressure [41]. The latter hypothesis that smooth muscle cells in the arterial wall seek axial steadiness but circumferential pulsatility.

#### 3.3.3. Opening Angle Analysis

The effect of the opening angle is illustrated in Fig. 10. By systematically varying the opening angle *β* from 0° to 120°, we assess how residual circumferential prestretch impacts the model’s mechanical predictions for absolute and relative outer diameters. The opening angle effect demonstrate the importance of the excised rings in the model workflow and its predictive influence.

**Figure 10:**
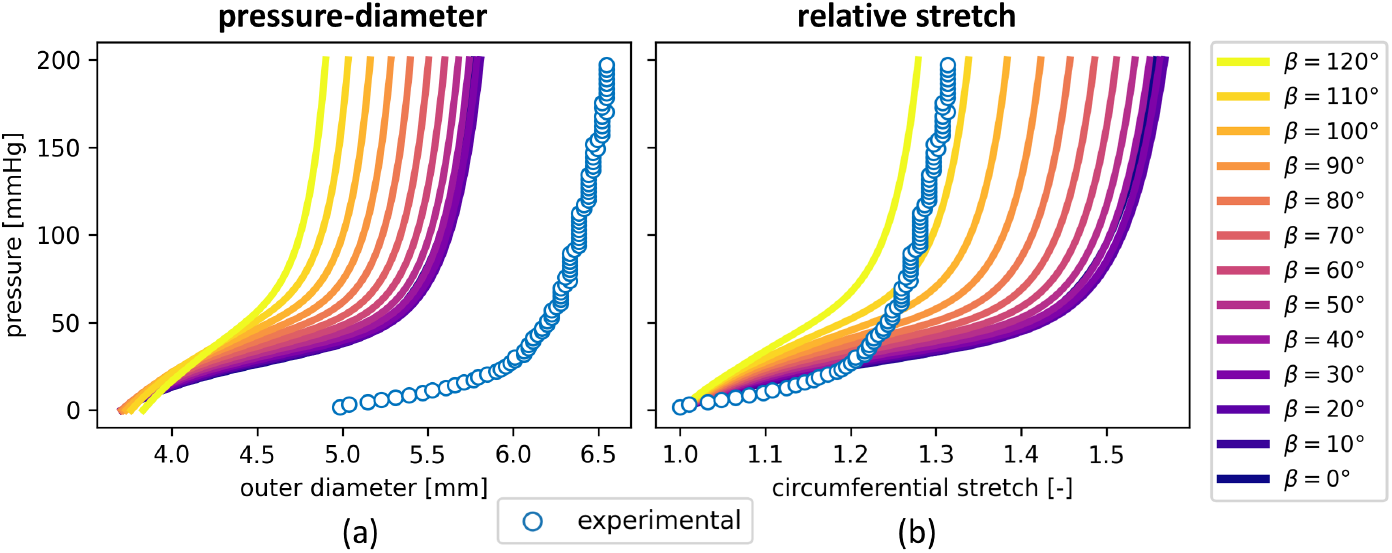
Opening Angle Sensitivity Analysis. Effect of a varying opening angle on model predictions for representative specimen B. The opening angle ranges from 0° to 120° in increments of 10°. (a) Modeled pressure–diameter curves for each opening angle value. (b) Corresponding relative circumferential stretch behavior. The experimentally measured opening angle is *β* = 77°.

##### Increasing the opening angle stiffens both the absolute and relative pressure–diameter curves

An increasing opening angle causes a stiffening of the pressure–diameter response, both in absolute values and in relative deformation (Figs. 10a and 10b). A larger opening angle implies greater residual stress at zero pressure, effectively pre-loading the vessel wall in the circumferential direction. As a result, the tissue deforms less under pressurization, leading to a stiffer mechanical response. For sample B, the experimentally measured opening angle was *β* = 77°, which aligns well with the model’s predicted behavior in this intermediate stiffness range. Importantly, the sensitivity of model predictions to variations in *β* suggests that incorporating accurate residual stress measurements is essential when simulating vascular mechanics. Unlike the tubular axial stretch, which is externally imposed during testing, the opening angle reflects intrinsic tissue architecture retrieved from ring opening experiments, and should thus carefully be considered during model calibration. Through Eq. (12) we perform an optimization of the optimal opening angle on the experimental extension–inflation data. Figure 7c retains the initial GOH parameters of Table 1 with *λ*_ax_ = 1.75, and finds the opening angle and thus circumferential prestretch that minimizes the relative outer diameter difference. This results in a value of *β* = 110.6°, leading to circumferential prestretches of *g*_circ,i_ = 1.09 and *g*_circ,i_ = 1.12, at the inner and outer wall, respectively. Our initial model was thus based on a underestimation of the opening angle. Note that the resulting circumferential prestretches are calculated through Eq. (4), and therefore highly dependent on the open and closed ring diameters, as well as the wall thickness *H*. An opening angle of *β* = 110.6° also leads to an updated axial prestretch *g*_ax_ = 0.78, ensuring a match in the experimental and modeled load-free thickness for *P* = 0 and *λ*_ax_ = 1.0.

### 3.4. Geometrical Uncertainty

In addition to the GOH material parameters, the axial stretch and circumferential prestretch, several other geometrical quantities remain uncertain across the different load-free and loaded configurations. These uncertainties are briefly discussed in the following section.

#### The measured load-free thickness and reference diameters are inconsistent in different configurations

Table 1 shows a discrepancy between the mounted outer diameter of the tubes, *D*_o_, and the measured outer diameter from the excised rings, 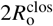. The mismatch could originate from the fact the the ring dimensions were measured at anatomical positions more distal or more proximal to the experimental tubes (Fig. 2), leading to sample-specific biological intravariability. A clinical validation could include a comparison between the *in vivo* arterial wall thicknesses and the modeled values of *h*, expressed in the deformed configuration, see Fig. 10. The unpressurized wall thickness used in the model, derived from the closed rings and defined as 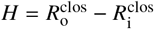, is also notably larger than the experimental thickness *T* measured from the squared biaxial sample (Table 1), which was used to calculate the planar-biaxial stress tensor 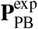. However, a thinner wall will intuitively result in even larger pressurized diameters, due to reduced resistance to circumferential expansion. Note the concave shape at low pressures remained because of the unaltered GOH parameters in this step.

#### Experimental pressure–diameter data should be enhanced with *in vivo* measurements

To obtain an exact and sample-specific axial stretch estimation, one should measure the arterial length *l* before explantation, and the excised length *L*^clos^ of the same tissue *ex vivo*, resulting in an estimation of *λ*_ax_. According to Fig. 3, the same axial measurement could be performed on the open ring *L*^open^, in order to obtain the experimental 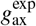. The experimental axial prestretch could then be compared to the modeled value in Table 1 to verify the workflow. However, in the current model workflow, the optimized *g*_ax_ guarantees computational consistency by ensuring a match between the experimental load-free closed ring configuration and the modeled geometry at *P* = 0 and *λ*_ax_ = 1.0. However, biological variability might surpass experimental accuracy when measuring the difference between *L*^clos^ and *L*^open^. An extra *in vivo* measurement could assess the diameter 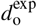 at zero pressure, considering the unpressurized artery before sacrifice. The model can predict the same 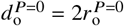 by solely applying the axial stretch *λ*_ax_ of the deformation gradient. Note that same the diameter at zero pressure after axial loading is also used to normalize the results and compute the relative circumferential stretch curves. Interestingly, still for 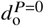 in Fig. 8a, we see that the experimental outer diameter from 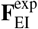 is close to the modeled 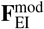at *λ*_ax_ = 1.0.

#### Experimental extension-inflation requires advanced imaging and can suffer from clamping boundary condition effects

The experimental extension-inflation test setup and resulting experimental pressure–diameter curve could be improved in order to enhance the accuracy and reliability of the measurements. For instance, a dual-camera system could be installed to capture uniform and out-of-plane three-dimensional deformations using 3D-DIC, and to track the actual outer diameter of the pressurized test sample, rather than relying on a two-dimensional plane projection. In addition, the experimentally applied extension and inflation conditions are susceptible to error. The inflation pressure may be affected by leakage or offsets in the pressure regulation system, and the actual axial stretch experienced by the sample may differ from the nominal value of *λ*_ax_ = 1.75 applied at the actuator level. The latter could be experimentally measured by marker tracking of the longitudinal axis extension. Pure pixel tracking of the sample height now shows *l* ^*P*=0^ /*L*^*^ = 1.74, see Table 1. During internal pressurization, the measured diameter may vary along the length, typically exhibiting a barrel-shaped profile. This length-dependent diameter evolution originates from the limited sample availability, where the available carotid samples should be free of holes or defects along the required testing length to ensure reliable data. In the current study, only the diameter at the middle of the sample is extracted from the extension-inflation experiments, and the model assumes a straight, perfect cylindrical geometry. Incorporating the length-dependent diameter evolution into the analysis could provide a more accurate description of the deformation. In the optimization described by Eq. (12), the inclusion of a longitudinal dimension could then be considered to capture length-dependent effects on the diameter evolution. On the modeling side, the behavior of a hollow thick-walled cylinder under internal pressure, axial stretch, and radial constraints could be examined. Such radial boundary conditions introduce longitudinal stress variations and shear stresses in the radial-axial plane, which can induce bulging or deflation of the cylinder and must be accounted for in a comprehensive model of extension-inflation behavior. Altogether, these results highlight the crucial interplay between accurate experimental measurements and endstage pressure–diameter results. While material calibration is essential, mismatches in the assumed wall thickness or vessel dimensions can propagate into model errors and obscure true mechanical behavior. This underscores the need for careful geometric validation in conjunction with mechanical testing for reliable computational modeling of vascular tissues. The total experimental and model uncertainty, together with the variability of the geometries and model parameters, are therefore also always important to consider. Literature presents different techniques to investigate this further [42–44].

## 4. Conclusions and Outlook

This study presents an integrated experimental and computational framework for characterizing ovine carotid artery biomechanics, bridging the gap between planar-biaxial testing and clinically relevant pressure–diameter behavior. We performed biofidelic extension-inflation experiments to compare model performance with benchtop pressure–diameter measurements. Our results highlight that translating laboratory measurements to realistic physiological conditions critically depends on accurately representing axial stretch and circumferential prestretch, as well as *in vivo* geometries. When these key parameters are well-characterized and their uncertainties properly accounted for, planar-biaxial stress– stretch data can become a reliable input for arterial pressure–diameter predictions.

We systematically combined planar-biaxial loading, ring opening experiments and extension-inflation tests on the same arterial tissue. Squared, ring and tubular samples of the same specimen allowed us to capture complementary loading conditions and tissue behaviors. To ensure physiological relevance, we designed tailored experimental protocols with controlled axial and circumferential loading in planar-biaxial tests, and axial loading under luminal pressure in extension-inflation setups. To minimize measurement uncertainty, we employed digital image correlation for accurate deformation tracking, and incorporated precise geometric data from micro laser scanning and ring-opening tests. These efforts improved the reliability of stress calculations and deformation measurements, thereby strengthening model input quality.

The modeling framework used the Gasser–Ogden–Holzapfel (GOH) constitutive model to capture the anisotropic hyperelastic material behavior of carotid arteries. Biomechanical predictions were performed through analytical thick-walled cylinder simulations with subject-specific tissue properties, boundary conditions and prestretches. Robust validation was achieved by benchmarking simulated pressure–diameter behavior against experimental extension-inflation data from the same specimens. Systematic comparison of absolute and relative pressure–stretch responses revealed key discrepancies and highlighted the influence of parameter choices on model performance.

Our results confirmed that the *in vivo* stretch in carotid arteries is higher in the axial direction, while the tissue exhibits lower circumferential prestretches. The GOH model fitted the planar-biaxial data well and proved suitable for pressure– diameter simulation. Comparisons between experimental and simulated pressure–diameter curves showed a systematic underestimation of the absolute pressure behavior and overestimation of the relative circumferential stretch, where we found that the anisotropic balance of the GOH model significantly shaped the pressure–diameter curve. We showed that the planar-biaxial stretch regime probes fundamentally different deformations compared to an extension-inflation test. We therefore fitted the GOH parameters directly to the pressure–diameter data, demonstrating the versatility of the constitutive parameters, depending on their biomechanical loading condition. Sensitivity analyses of the tubular axial stretches and ring opening angles emphasized their importance in accurate model calibration. while an increased axial stretch reduced the predicted absolute diameters, it increased the relative outer diameters. An increasing opening angle led to an overall stiffening of the pressure–diameter behavior. Optimizing the axial stretch and circumferential prestretch further enhanced the biofidelity of our simulations. Finally, we showed how axial loading is critical when performing parameter optimizations based on extension-inflation data.

Despite the robustness of our approach, some limitations remain. Experimental noise, biological heterogeneity, and model simplifications contribute to uncertainty. Standardizing measurement protocols, improving accuracy in thickness and diameter measurements, and automating post-processing workflows will help reduce variability. Future work should focus on (I) expanding datasets to quantify aleatoric and epistemic uncertainties, (II) incorporating clinical imaging data for further validation, (III) exploring 3D optical measurements and inverse modeling to refine experimental and modeling assumptions, (IV) applying constitutive artificial neural networks to autonomously discover novel strain energy density functions for carotid arteries, and (V) implement length-dependent diameter evolutions in the model, in order to consider longitudinal stress variations due to boundary condition effects.

## Data Availability

All source data, processing code and results are available in KU Leuven’s research data repository. The full dataset and associated materials will be made openly accessible upon journal acceptance.

## Acknowledgements

This work was supported by the doctoral fellowship SB1SE2125N from the Research Foundation Flanders (FWO) to Thibault Vervenne, by a doctoral fellowship 1157325N from the FWO to Nele Demeersseman, and by the research project G029819N from the FWO to Nele Famaey. The authors would also like to extend their gratitude to Kimberly Crevits, lab manager at FIBEr, for her invaluable support in enabling high-quality mechanical testing of biological tissues at KU Leuven.

## Appendices

**Table A1:** Biaxial Dimensions and Fitted GOH Material Parameters. Biaxial square dimensions and Gasser–Ogden–Holzapfel (GOH) material parameters θ = {*c*_1_, *k*_1_, *k*_2_, *κ, α*}, fitted to the fifth loading cycle of biaxial tensile tests for samples A to E. The values *W*_1_ and *W*_2_ are the perpendicular widths of the axial and circumferential direction, respectively. *T* is the averaged wall thickness of the biaxial square sample measured through micro laser scanning (MLS). The goodness of fit of the modeled and experimental stress–stretch curves are highlighted by *R*^2^.

| sample | squares [mm] | | | GOH parameters | | | | | $R^2$ [-] |
| --- | --- | --- | --- | --- | --- | --- | --- | --- | --- |
| | $W_1$ | $W_2$ | $T$ | $c_1$ [kPa] | $k_1$ [kPa] | $k_2$ [-] | $\kappa$ [-] | $\alpha$ [rad] | |
| A | 10.5 | 10.6 | 0.43 | 28.4 | 7.0 | 6.259 | 0.112 | 0.000 | 0.874 |
| B | 9.9 | 10.8 | 0.30 | 49.2 | 5.3 | 9.318 | 0.078 | 0.000 | 0.908 |
| C | 10.9 | 9.7 | 0.23 | 67.2 | 3.0 | 12.268 | 0.134 | 0.000 | 0.796 |
| D | 10.3 | 10.7 | 0.31 | 31.2 | 18.9 | 6.469 | 0.186 | 0.000 | 0.853 |
| E | 10.7 | 10.9 | 0.57 | 20.8 | 7.8 | 6.595 | 0.136 | 0.000 | 0.883 |

**Table A2:** Dimensional Measurements and Prestretch Variables. Tube dimensions and ring measurements for samples A to E. *L* represents the harvested axial length of the tubes. *D*_o_ highlights the mounted and unpressurized outer diameters, while 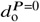 describes the measured unpressurized outer diameters after applying the axial stretch 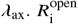 and 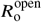 are the inner and outer radii of the ring in the open configuration, while 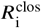 and 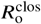 are the corresponding radii in the closed configuration. *β* is the opening angle in degrees, *g*_circ,i_ and *g*_circ,o_ are the resulting inner and outer circumferential prestretches respectively, and *g*_ax_ is the optimized axial prestretch.

| sample | tubes [mm] |  |  | rings [mm] |  |  |  | [°] | [-] | [-] | [-] |
| --- | --- | --- | --- | --- | --- | --- | --- | --- | --- | --- | --- |
| | $L$ | $D_o$ | $d_o^{P=0}$ | $R_i^{\text{open}}$ | $R_o^{\text{open}}$ | $R_o^{\text{clos}}$ | $R_i^{\text{clos}}$ | $\beta$ | $g_{\text{circ},i}$ | $g_{\text{circ},o}$ | $g_{\text{ax}}$ |
| A | 36 | 5.36 | 4.84 | 1.95 | 3.10 | 2.58 | 1.85 | 44 | 1.08 | 0.95 | 0.98 |
| B | 52 | 5.49 | 4.99 | 2.40 | 3.15 | 2.45 | 1.83 | 77 | 0.97 | 0.99 | 1.04 |
| C | 54 | 6.18 | 5.59 | 2.60 | 3.15 | 2.63 | 2.08 | 42 | 0.90 | 0.94 | 1.17 |
| D | 30 | 6.04 | 5.47 | 2.85 | 3.70 | 2.80 | 2.00 | 94 | 0.95 | 1.02 | 1.02 |
| E | 49 | 4.82 | 4.42 | 3.30 | 3.90 | 2.53 | 1.85 | 131 | 0.88 | 1.02 | 1.11 |

**Figure A1:**
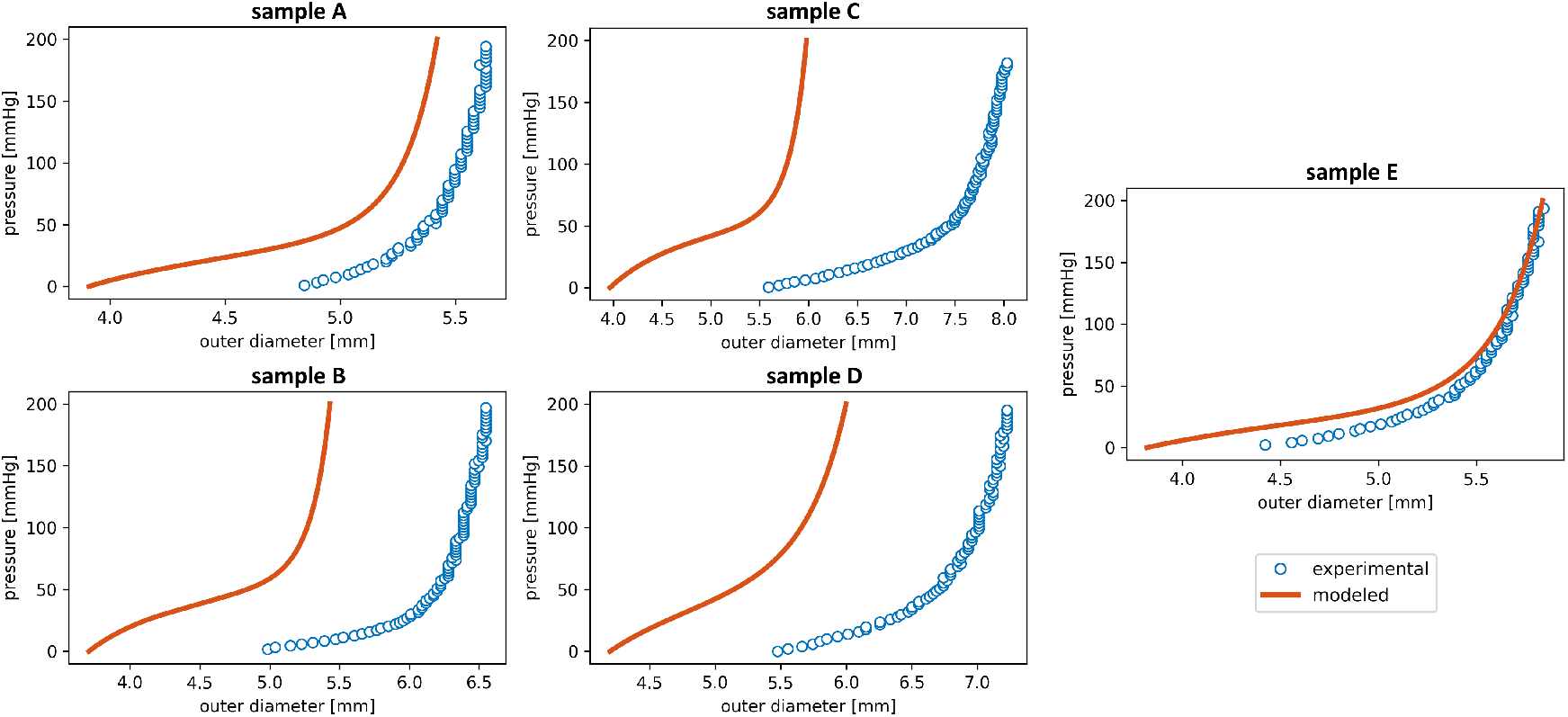
Modeled and Experimental Absolute Pressure–Diameter Behavior. Comparison of modeled and experimental pressure– diameter (*P*–*d*) relationships for the five carotid artery specimens (A–E). The modeled curves are generated using GOH material parameters fitted from the corresponding biaxial stress–stretch data. Experimental pressure–diameter data is extracted from extension-inflation tests performed on each specimen.

**Figure A2:**
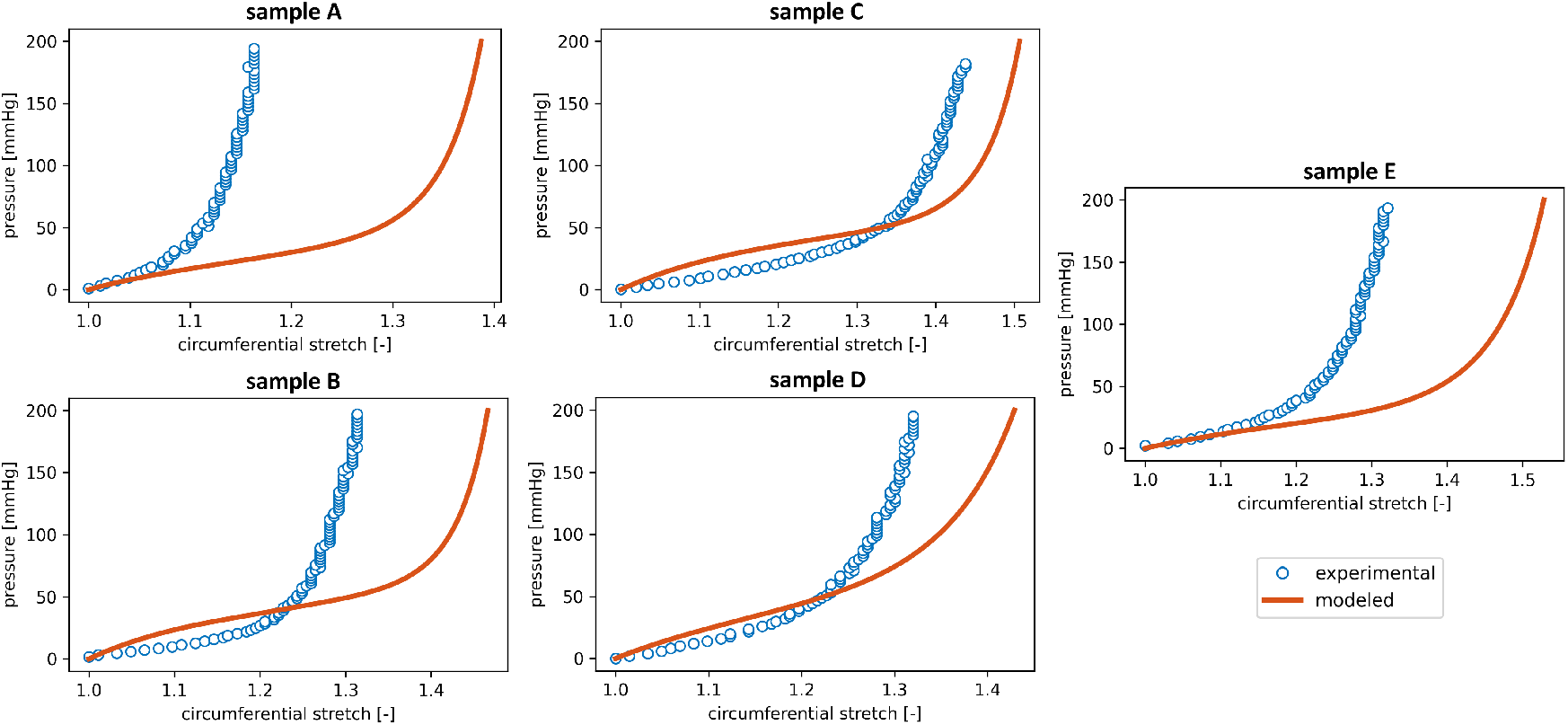
Modeled and Experimental Relative Pressure–Diameter Behavior. Comparison of modeled and experimental pressure– circumferential stretch (*P*–*λ*_circ_) relationships for the five carotid artery specimens (A–E).

**Figure A3:**
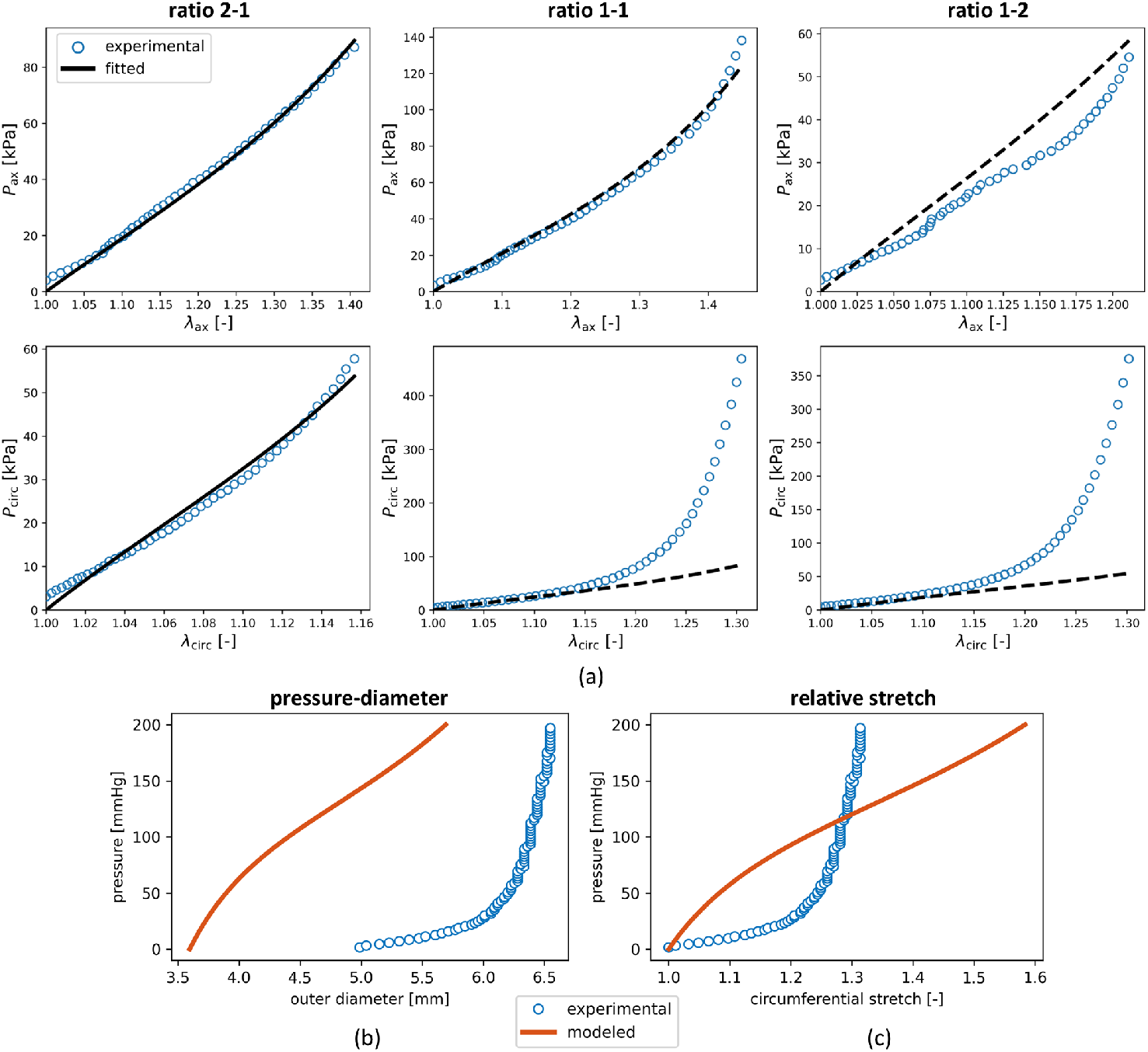
GOH Fitting and Model Predictions for the 2:1 Loading Ratio. Model results when the material parameters of the GOH constitutive model are fitted solely using the planar-biaxial data from the 2:1 axial-to-circumferential stretch ratio. The resulting material parameters for sample B are *c*_1_ = 35.2 kPa, *k*_1_ = 8.2 kPa, *k*_2_ = 5.5086, *κ* = 0.000 and *α* = 0.2761 rad. (a) Axial and circumferential stresses plotted under three loading protocols. Experimental stress–stretch data are shown alongside the fitted and predicted stress responses. (b) Resulting modeled and experimental absolute pressure–diameter behavior. (c) Corresponding pressure–circumferential stretch curves, showing relative deformation.

**Figure A4:**
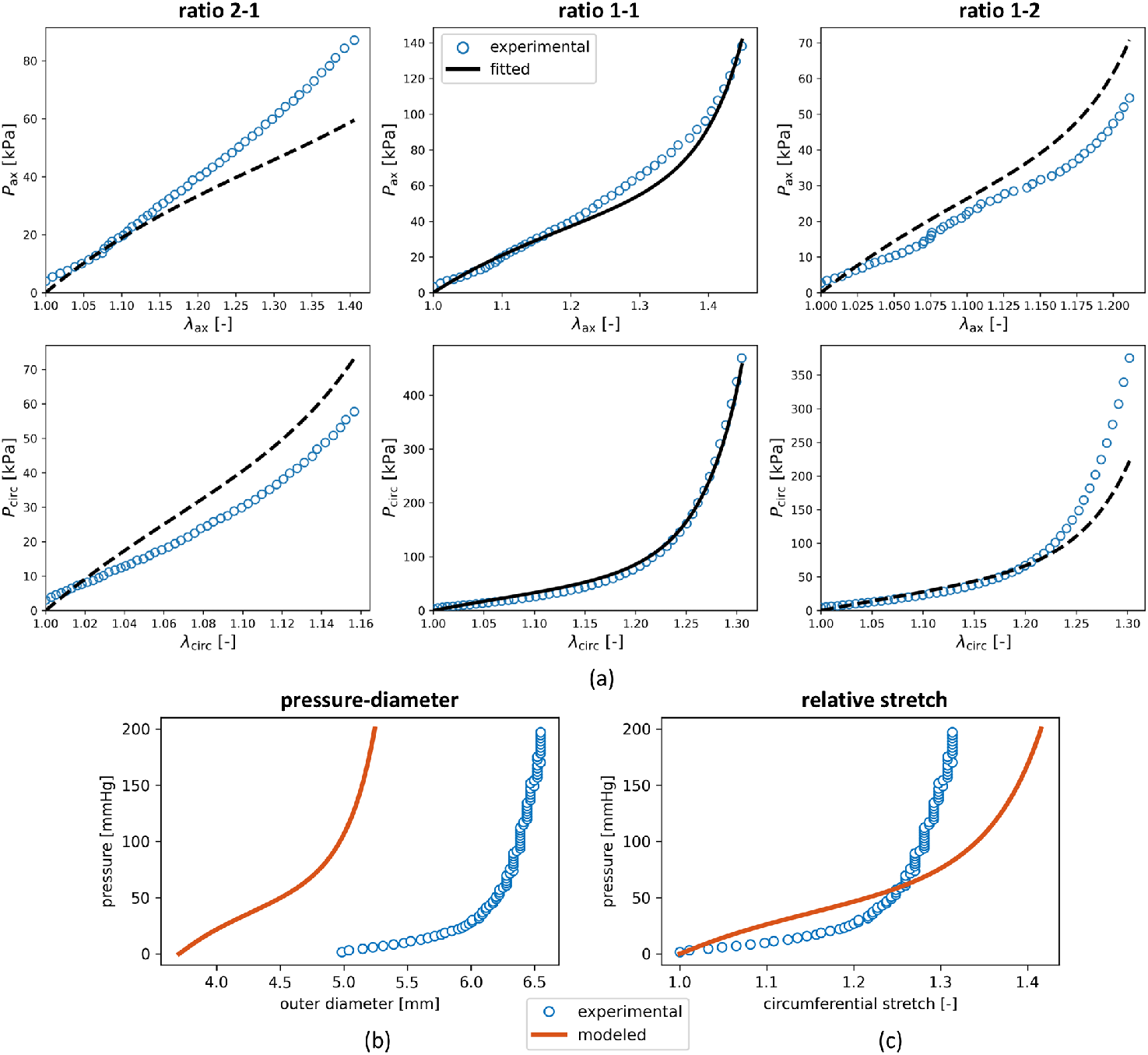
GOH Fitting and Model Predictions for the 1:1 Loading Ratio. Model results when the material parameters of the GOH constitutive model are fitted solely using the planar-biaxial data from the 1:1 axial-to-circumferential stretch ratio. The resulting material parameters for sample B are *c*_1_ = 45.2 kPa, *k*_1_ = 8.9 kPa, *k*_2_ = 4.4479, *κ* = 0.000 and *α* = 0.4088 rad. (a) Axial and circumferential stresses plotted for sample B under three loading protocols. Experimental stress–stretch data are shown alongside the fitted and predicted stress responses. (b) Resulting modeled and experimental absolute pressure–diameter behavior. (c) Corresponding pressure–circumferential stretch curves, showing relative deformation.

**Figure A5:**
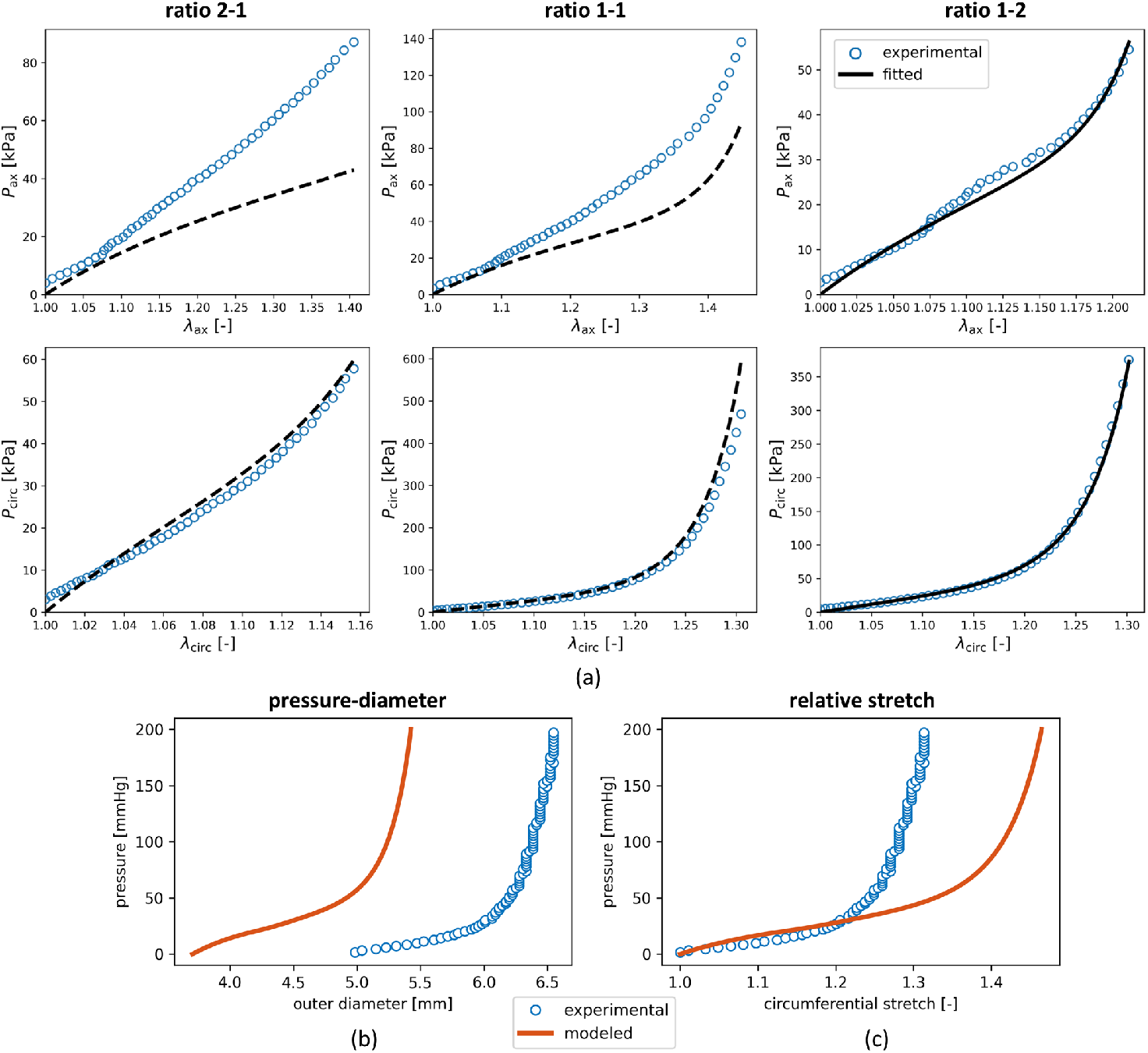
GOH Fitting and Model Predictions for the 1:2 Loading Ratio. Model results when the material parameters of the GOH constitutive model are fitted solely using the planar-biaxial data from the 1:2 axial-to-circumferential stretch ratio. The resulting material parameters for sample B are *c*_1_ = 37.4 kPa, *k*_1_ = 43.8 kPa, *k*_2_ = 0.8815, *κ* = 0.2358 and *α* = 1.0515 rad. (a) Axial and circumferential stresses plotted under three loading protocols. Experimental stress–stretch data are shown alongside the fitted and predicted stress responses. (b) Resulting modeled and experimental absolute pressure–diameter behavior. (c) Corresponding pressure–circumferential stretch curves, showing relative deformation.

**Figure A6:**
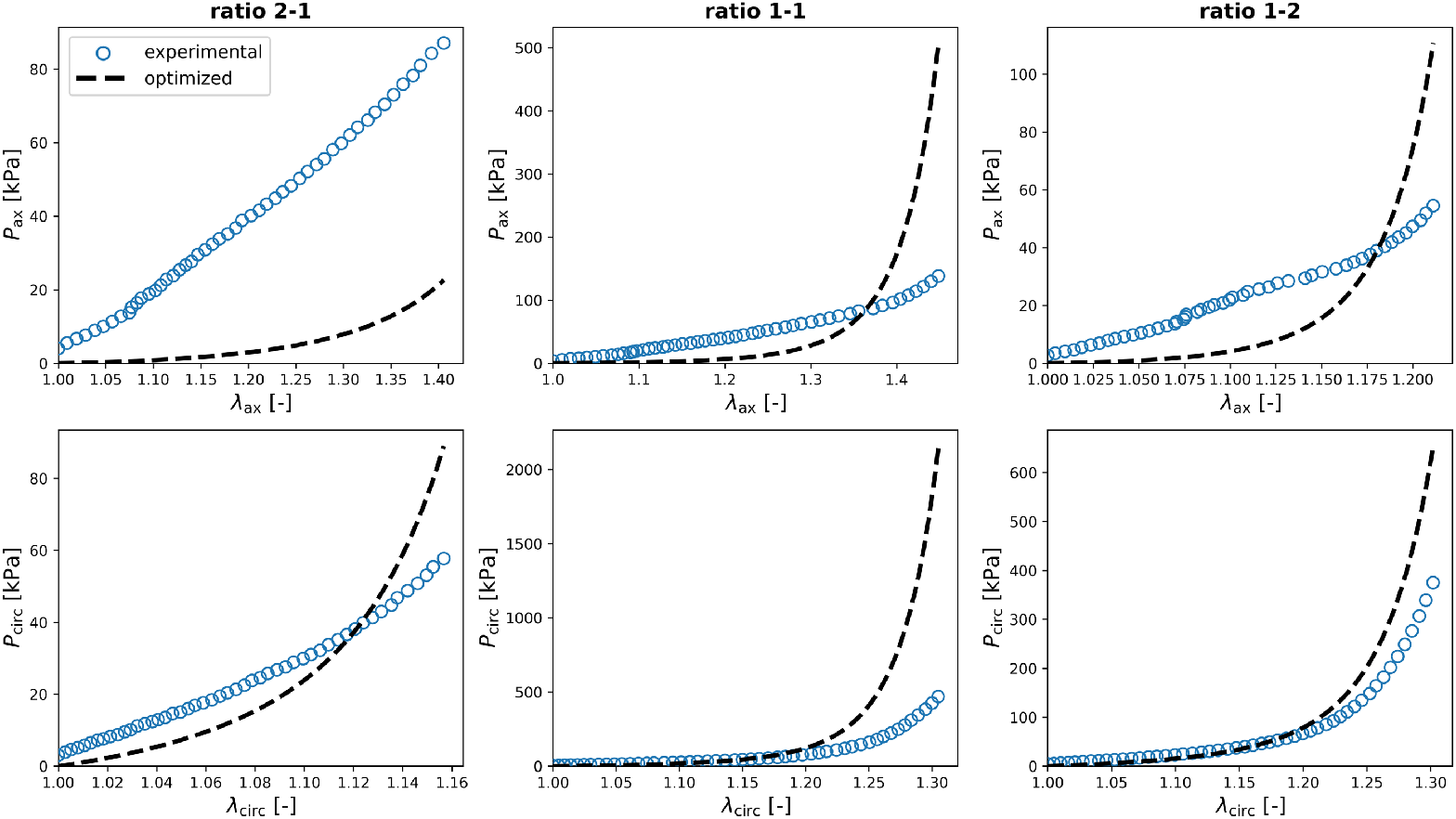
Experimental and Virtual Planar-Biaxial Stress–Stretch Curves. Axial and circumferential first Piola-Kirchhoff stresses (*P*_ax_ and *P*_circ_) plotted for sample B under three loading protocols with stretch ratios of 2:1, 1:1, and 1:2 in the axial and circumferential directions (*λ*_ax_ and *λ*_circ_). Experimental stress–stretch data are shown alongside the the virtual stress responses computed using the optimized material parameters of the GOH constitutive model (*c*_1_ = 0.632 kPa, *k*_1_ = 39.5 kPa, *k*_2_ = 11.38, *κ* = 0.156 and *α* = 0.087 rad). These model parameters were obtained by fitting the GOH model directly to the pressure–diameter data from the experimental extension-inflation test (see Section 3.3.1 and Fig. 7a).

**Figure A7:**
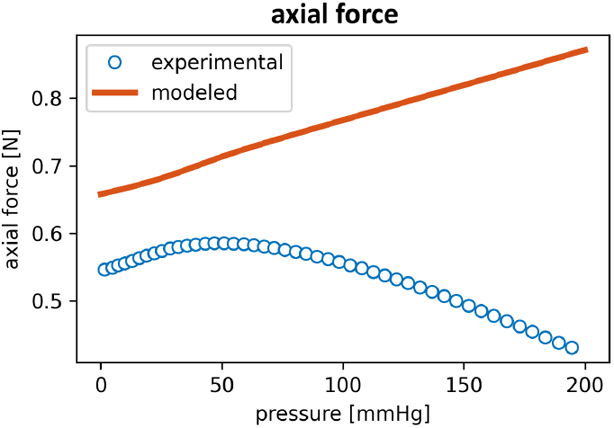
Experimental and Modeled Axial Forces. For the physiological pressure range of representative sample B, and at an axial stretch of *λ*_ax_ = 1.75, the experimentally measured axial forces during extension-inflation are plotted against the modeled values of *f*_ax_.

### Constitutive Artificial Neural Networks

Instead of relying on a predefined strain energy density function, we explore automated model discovery by training a constitutive artificial neural network (CANN). In this approach, we extend the set of strain invariants to include the isotropic second invariant 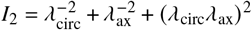 and the anisotropic fifth invariant 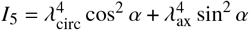. In the first layer of the neural network, we allow both the invariants and their squares, (·) and (·)2, as inputs. The second layer applies the identity function (·) and exponential function exp(·) to these transformed inputs. The layers are interconnected by carefully designed activation functions that ensure thermodynamic consistency, material objectivity, symmetry, physical restrictions, and polyconvexity [31]. Together, the neural network architecture gives rise to the following generalized strain energy density function:

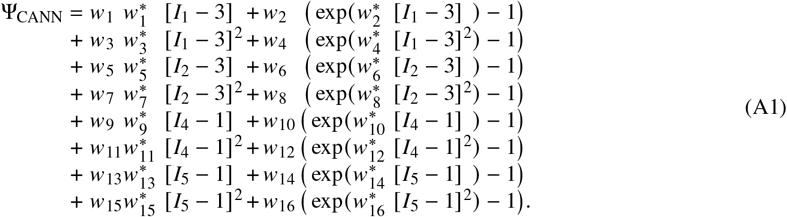

The network has a total of two times 16 trainable weights, resulting in more than 65,000 possible combinations. The network can learn any strain energy function constructed from first- and second-order, as well as identity and exponential additive combinations of the invariants *I*_1_, *I*_2_, *I*_4_, and/or *I*_5_. We also fit the fiber angle *α*, influencing the contribution of both the anisotropic terms *I*_4_ and *I*_5_. More details on the network’s architecture can be found in prior works [32, 34, 45].

Here, we use cross-sample feature selection to setup the loss function of Ψ_CANN_ (*I*_1_, *I*_2_, *I*_4_, *I*_5_). We train the neural network on all planar-biaxial samples together, with the fifth loading cycle of the three stretch ratios. We train for 8,000 epochs, with a batch size of 32. Early stopping is allowed within 2,000 epochs of no accuracy change [46, 47]. The adaptive Adam algorithm is used for first-order optimization. To prevent local minima effects, we initialize the weights of the neural network layers with random values drawn from a uniform distribution. Specifically, the Glorot normal initializer is used for the identity functions, and an unseeded random uniform initialization is used for the exponential functions, with a minimum and maximum weight value of 0.0001 and 0.1 respectively [32].

We enforce the weights to always remain non-negative, 𝒲 ≥ 0. L1 or Lasso regularization can induce additional sparsity by reducing some weights exactly to zero, which effectively reduces model complexity and improves interpretability [48]. Here, we set the parameter *α*_reg_ to 0.01 to activate regularization.

With our set of weights 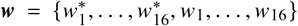 we perform a gradient descent learning on a weighted least-squares error loss function *L*, penalizing the error between the discovered planar-biaxial model 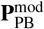 and the experimental planar-biaxial data 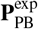

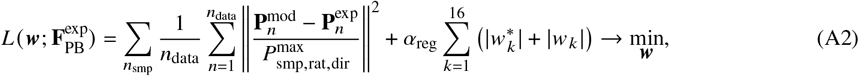

where *n*_data_ is the total number of data points of a specific sample, considering the three axial-vs-circumferential experimental loading ratios {1:1, 2:1, 1:2}, and the two loading directions: axial and circumferential, {ax, circ}. We extend the loss function with *n*_smp_ to account for the sample set from the different planar-biaxial tests. To account for all experiments equally, we weight the error with the maximum stress per ratio and per direction and for each sample,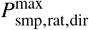. This loss function allows us to discover a unique model or set of invariant features and find a universal constitutive behavior. In the absence of radial stresses during the planar-biaxial test, we define an explicit expression for the Lagrange multiplier *p* as boundary condition for the incompressibility requirement.

Figure A8a demonstrates the CANN’s ability to identify physically interpretable and directionally dependent components of arterial tissue mechanics. The learned strain energy function includes the linear isotropic contribution from the first invariant in the form of [*I*_1_ − 3], with weights *w*_1_ = 90.034 kPa and 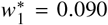, as well as the quadratic isotropic contribution of the first invariant [*I*_1_ − 3]^2^, with weights 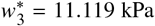 kPa and *w*_3_ = 0.111. An anisotropic exponential quadratic term exp([*I*_5_ − 1]^2^) is aligned with the circumferential direction (collagen fiber angle *α* = 0) and scales with weights *w*_15_ = 2.373 kPa and 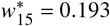. The goodness of fit across samples can be assessed through the mean experimental curve, or by aggregating the planar-biaxial data of all *n* = 5 samples for the three ratios and two directions, resulting in an *R*^2^ = 0.631, calculated w.r.t. the CANN-modeled stress–stretch curves 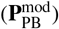. Intuitively, *R*^2^ is lower for cross-sample assessment, but previous work has shown that the neural network generally outperforms the GOH model [34]. The corresponding CANN weights indicate the relative importance of each term and reflect meaningful mechanical behavior, learned directly from the data. For carotid arteries, Eq. (A1) thus reduces to following strain energy density function:

**Figure A8:**
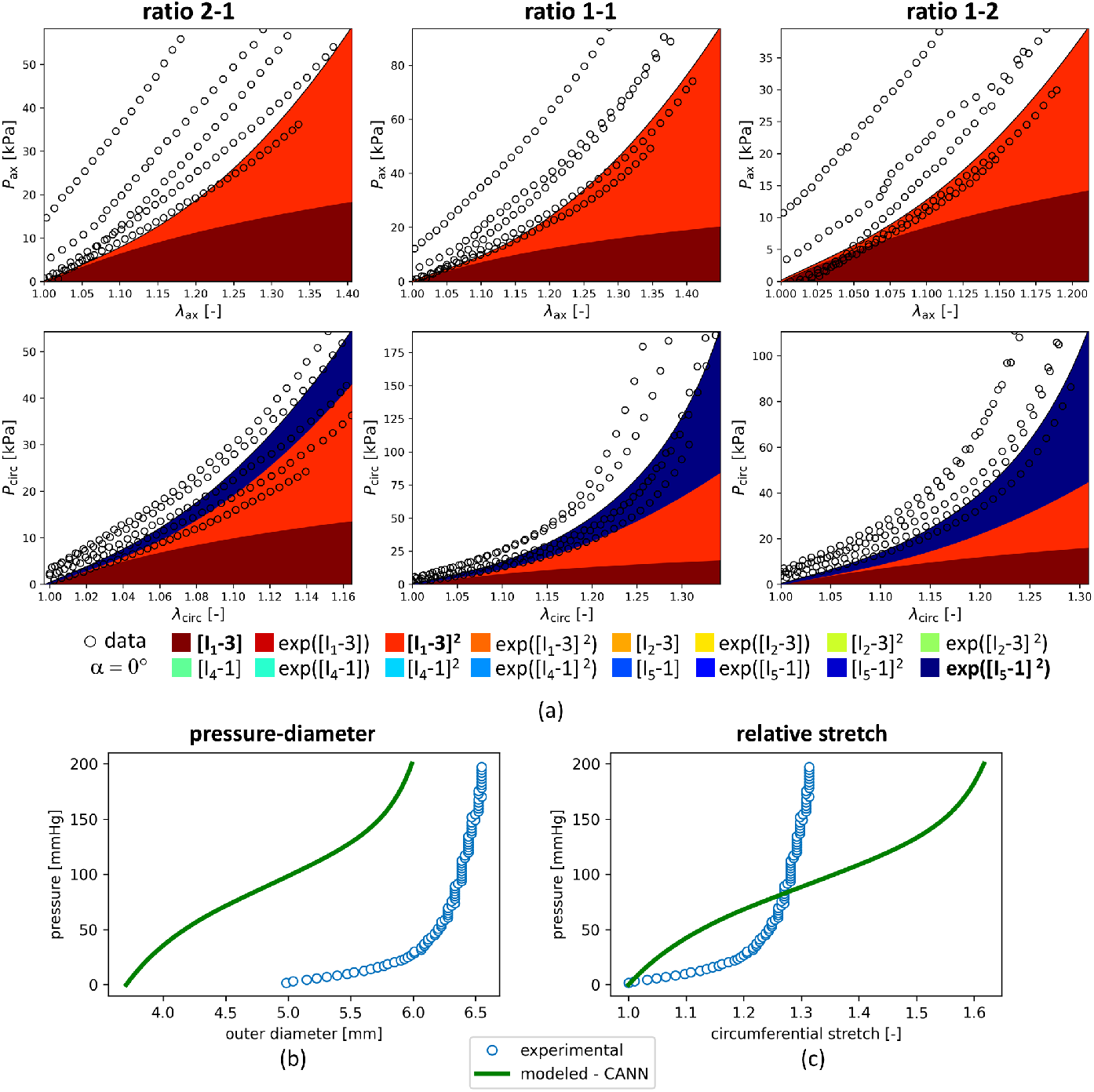
Automated Constitutive Model Discovery and Resulting Predictions. A constitutive artificial neural network (CANN) is trained on the planar-biaxial stress–stretch data from the five carotid artery specimens combined, and used in the model to predict pressure– diameter behavior. (a) The discovered strain energy function includes isotropic contributions from the first invariant in the form of [*I*_1_ - 3] and [*I*_1_ - 3]^2^, and a circumferential anisotropic fiber contribution represented by exp [*I*_5_ - 1]^2^). (b) Comparison of modeled and experimental absolute pressure–diameter behavior, where the discovered terms and learned weights are used to simulate the arterial wall response. (c) Corresponding pressure–circumferential stretch curves, showing relative deformation.

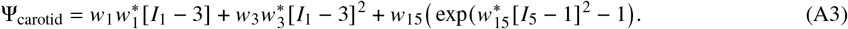

Figures A8b and A8c apply the discovered model to specimen B within the modeling framework. Using the learned invariant terms and weights, the model more accurately reproduces the absolute pressure–diameter response, and closely matches the experimental outer diameters at diastolic and systolic carotid blood pressures. While the relative pressure–circumferential stretch behavior captures the general trend, it tends to overshoot more noticeably in the high-pressure regime. This can be attributed to the dominant contribution of the anisotropic exponential *I*_5_ term, which is oriented solely in the circumferential direction and has a major influence in the physiological high-pressure regime of carotid arteries. In contrast, the isotropic *I*_1_ term induces a concave mechanical behavior primarily in the low-pressure regime, capturing initial vessel response. Through automated model discovery, these result highlights the CANN’s ability to uncover interpretable and predictive constitutive forms without predefined assumptions. It offers an attractive, data-driven pathway to model development in pressure–diameter simulations based on planar-biaxial datasets. We show that the constitutive neural network robustly discovers similar *I*_1_ and *I*_5_ invariant forms of arterial constitutive models for carotid arteries as observed in other pressurized cardiovascular tissues, such as human aortic arches or ovine main pulmonary arteries [34, 45].

## Notes

### Competing Interest Statement

The authors have declared no competing interest.

### Summary of Updates

upload of the revised manuscript with adapted appendices

